# Combinatorial phosphorylation on CTD of RNA polymerase II selectively controls transcription and export of protein-coding mRNAs

**DOI:** 10.1101/2025.01.13.632769

**Authors:** Svetlana B. Panina, Seema Irani, Haley A. Hardtke, Renee Stephenson, Brendan M. Floyd, Rosamaria Y. Moreno, Edward M. Marcotte, Qian Zhang, Y. Jessie Zhang

**Affiliations:** Department of Molecular Biosciences, The University of Texas at Austin, 2500 Speedway, Austin, TX, USA

## Abstract

C-terminal domain (CTD) of RNA polymerase II is crucial for recruiting transcription regulators via specific post-translational modifications (PTM), especially phosphorylation. The hypothesis of combination of PTMs, or ‘CTD code’, that can allow precise and dynamic recruitment of transcription machinery is highly attractive, yet the experimental evidence to support this hypothesis has been scarce. Here, despite lacking specific antibodies for combinatorial CTD phosphorylation, we developed an innovative approach that detects double phosphorylation patterns on the CTD in a whole-genomic fashion by leveraging the antibody masking effect with selectively removing the flanking interference. Using this method, we detected pT4pS5 double phosphosites occurring exclusively during the transcription of protein-coding genes. Furthermore, we showed that pT4pS5 marks recruit the Transcription and Export complex (TREX), which specifically facilitates mRNA processing and nucleocytoplasmic export of protein-coding mRNAs. The recruitment of TREX by pT4pS5 phosphosites is particularly important for the processing of lengthy neurogenesis-related genes. Our results provide experimental support for the notion that CTD coding system can function combinatorially and in a gene-specific manner, which encodes an exact information about the transcription of specific gene clusters. This method can be broadly applied to map all combinatorial PTM patterns on RNA polymerase II, paving the way for a deeper understanding of gene-specific transcription regulation at the molecular level.

## INTRODUCTION

The C-terminal domain (CTD) of the largest subunit of RNA polymerase II (Pol II), Rpb1, is a unique domain essential for all Pol II-mediated transcription including all protein-coding mRNAs and some noncoding RNAs in eukaryotes (*1*). The CTD facilitates the recruitment of various transcriptional regulatory proteins spatiotemporally, which are crucial for progressing through transcription (*2*). The loss of the CTD is lethal for cells, with at least about one-third of its repeats required for cell survival (*2*). The CTD functions as a hub for transcriptional regulation through its extensive post-translational modifications (PTMs), with phosphorylation being the most prominent during active transcription (*3*). Despite its simplicity - composed of tandem repeats of a consensus heptad sequence (Y^1^S^2^P^3^T^4^S^5^P^6^S^7^) - five residues out of the heptad are subject to phosphorylation, while the two proline residues undergo isomerization, influencing the recognition of phosphorylation patterns (*4*).

This potential to combine different PTM states within the heptad or neighboring repeats makes the CTD an intricate regulatory platform (*5*). Recent studies challenge the traditional view of the CTD as a passive scaffold for protein recruitment, but instead, suggest a model that the CTD actively encodes transcriptional outcomes by creating combinatorial PTM patterns in response to developmental and environmental stimuli (*6*). This dynamic system allows transcription regulators to be recruited to specific PTM configurations, ultimately directing different transcriptional outcomes. In this model, Pol II functions as both the recipient and the integrator of PTM signals, responding to chromatin states and external cues such as nutrient availability, stress, or DNA damage (*7–9*). This coupling is critical for regulating gene expression programs under varying physiological conditions. In such context, combinatorial PTM patterns have high power in encoding information with precision. Supporting this notion, mass spectrometry analysis of human and yeast CTDs has identified the occurrence of double phosphorylation within heptad repeats (*10, 11*). However, experimental evidence linking these combinatorial phosphorylation patterns to specific regulatory functions on Pol II remains limited without knowing which stage of transcription such PTMs occur.

The bottleneck to study the combinatorial phosphorylation in CTD is the absence of highly specific antibodies capable of recognizing such patterns in cells (*6*). The ideal antibody would bind specifically to double phosphorylation events while showing minimal cross-reactivity with individual phosphorylations or other modifications. With all different double phosphorylation patterns identified by mass spectrometry in human cells (*10*), few antibodies targeting double-phosphorylated CTD motifs are available, preventing comprehensive mapping of these modifications during transcription. The inability to pinpoint the genomic locations of specific double phosphorylation marks on the CTD severely limits our understanding of their functional roles (*12*). Although proteomic studies have identified hundreds of proteins binding to phosphorylated CTD residues, the specific epitopes recognized remain poorly defined. Significant overlap exists among the proteins interacting with single phosphorylation sites, suggesting that multiple phosphorylation patterns within the CTD could provide the necessary specificity for transcriptional regulation. It is essential to identify PTM patterns during transcription and identify the CTD-binding transcription regulators they recruit to modulate transcription.

To overcome this challenge, we took advantage of our previous observation that some CTD antibodies can be masked by nearby phosphorylation (*13*). In this study, we explored the possibility of removing such interfering phosphorylation specifically by phosphatases, thereby potentially identifying the genomic locations where the two phosphorylations co-occur. Hence, we took advantage of a highly selective phosphatase Ssu72 to specifically remove the flanking Ser5 phosphorylation, which had been blocking the recognition of Thr4 by the phospho-Thr4-specific antibody 6D7. A pairwise comparison of ChIP-seq profiles, with and without Ssu72 phosphatase activity, allowed us to map the locations of pT4pS5 double phosphorylation by detecting the unmasked phospho-Thr4 peaks. Surprisingly, we found that pT4pS5 double phosphorylation is absent in small regulatory RNAs transcribed by Pol II but is only present in protein-coding genes, both at transcription start sites (TSS) and within gene bodies. Proteomic analyses further identified the TRanscription and EXport complex (TREX) among binding partners recruited by pT4pS5 double phosphorylation marks, with cellular experiments validating this association. Consistent with the function of TREX in transcript export and splicing, the loss of pT4pS5 marks, especially intronic-mapped, leads to alternative selection of splicing sites, especially in long genes important for neurogenesis and neurodevelopment. Critically, the nucleocytoplasmic transport of specific mature mRNAs is hampered upon the loss of pT4pS5 double phosphorylation. Overall, our study reveals that the pT4pS5 double phosphorylation is a gene-specific CTD code for protein-coding genes that recruits specific regulators such as TREX for transcript export out of the nucleus and mRNA maturation.

## RESULTS

### A strategy to map the genomic locations of double phosphorylation patterns on the CTD

Mass spectrometric studies in human cells detected that pT4 phospho-mark frequently co-occurred with pSer2 and pSer5 within the same CTD heptad, and pT4pS5 double phosphorylation pattern was prevalent in both yeast and human cells, accounting for 15-20% of all double-phosphosites (*10*). Yet, exact genomic location and functional properties of such double-marks in the transcription cycle remain unknown. In our recent study investigating the whole-genome pattern of Thr4 phosphorylation on the CTD, we uncovered under-appreciated occurrence of pT4 playing role in pausing release and 3’ end processing, previously undetected by the antibody 6D7 when masked by nearby phosphorylation (*13*). The presence of significant masking effect indicates that Thr4 phosphorylation is frequently flanked by nearby phosphorylation on CTD.

Hence, we pondered the possibility of mapping the genomic location of pT4pS5 combinatorial double-phosphosites within the same CTD heptads by comparing the pT4 signal with or without unmasking. We can achieve that via differential binding analysis between samples treated with highly specific phosphatase Ssu72 and catalytically inactive version of the enzyme (mutant). In this case, pT4 ChIP-Seq peaks appearing when compared to samples without phosphatase activity treatment indicate the position where the double phosphorylation occurs (**Fig. 1A**).

**Figure 1.**
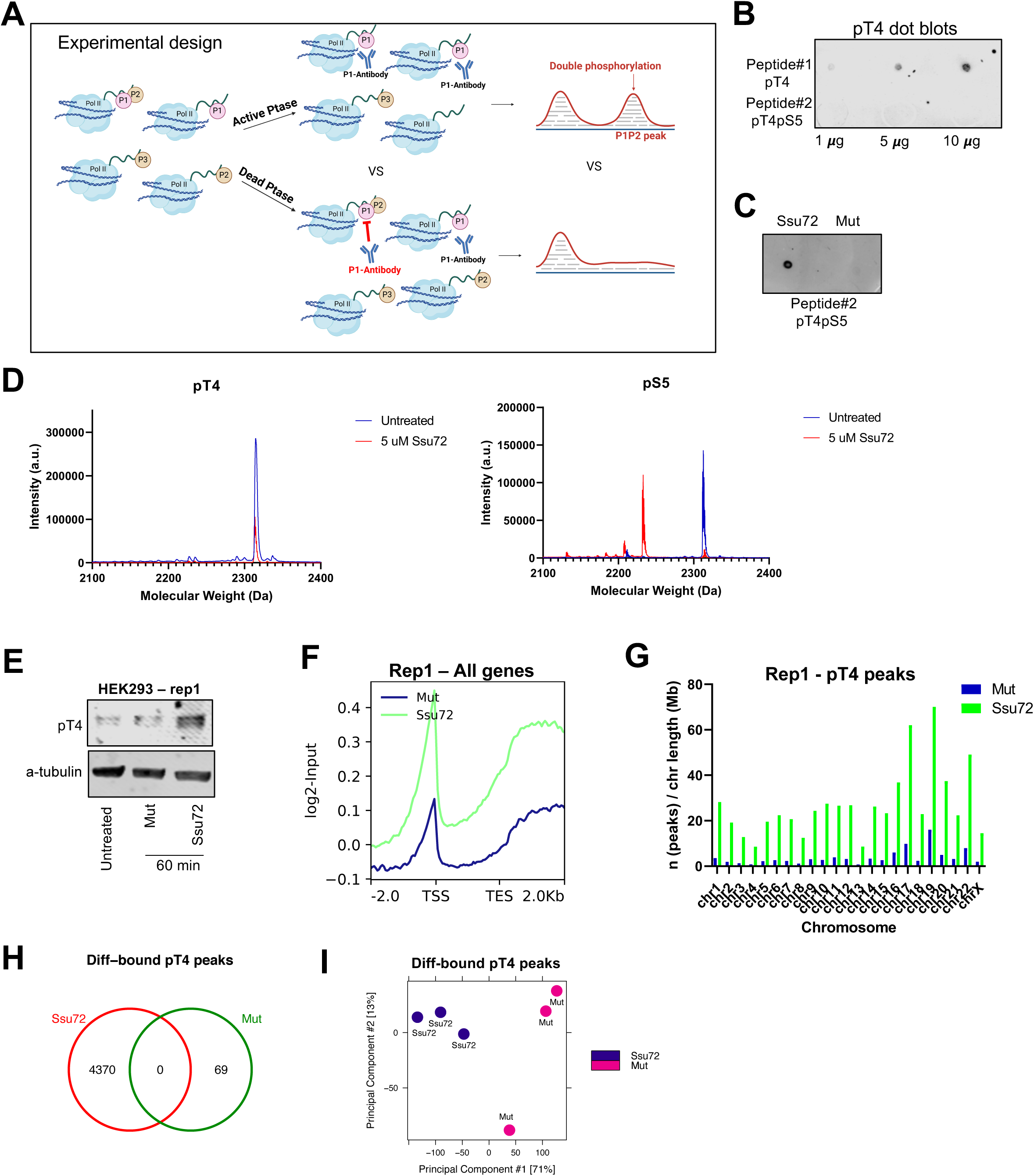
A framework for detection of combinatorial CTD modifications using a pair of active and inactive phosphatases. **A,** Experimental and analytical framework of detection of double phosphorylation sites (P1P2). The schematic was created using BioRender software. **B,** Dot blot performed using pT4 antibody (6D7) and serial dilutions of synthetic peptides with singly-(pT4, #1) and doubly-phosphorylated (pT4pS5, #2) residues. **C,** Dot blot performed using pT4 antibody (6D7) and spotted doubly-phosphorylated peptide (pT4pS5) after incubation with active Ssu72 phosphatase or inactive phosphatase (Mut) for 30 min. One representative replicate is shown. **D,** MALDI-TOF profiles of untreated (in blue) synthetic peptides having pT4 mark (*left*) or pS5 mark (*right*) vs. peptides treated with 5 µM of active Ssu72 phosphatase (in red). **E,** Western blot demonstrating significant increase in pT4 signal upon treatment of cellular lysate (HEK293 cells) with active Ssu72 phosphatase for 60 min. One representative replicate is shown. **F,** Whole-genome metagene (n = 60,264) profile of Input-normalized pT4 ChIP-Seq signal under Ssu72 vs Mutant treatment. Region between TSS (transcription start site) and TES (transcription end site) is scaled to 2,000 bp. Every bin is 50 bp. One representative replicate is shown. **G,** pT4 peak distribution by chromosomes normalized by every chromosome’s length in Mb upon Ssu72 vs Mutant treatment. One representative replicate is shown. **H,** Venn diagram showing absolute numbers of gained or lost pT4 peaks in Ssu72 vs Mut condition called by ‘DiffBind’ package in R. **I,** PCA analysis of differentially bound pT4 peaks in Ssu72 vs Mut condition.

For this strategy to work, we first need to identify a nearby modification that blocks the antibody for antigen (pT4) recognition. Phospho-Thr4 antibody 6D7 has been widely used but its recognition can be blocked by nearby Ser5 and Ser2 phosphorylation (*14*). To validate that, we confirmed that pT4 antibody (6D7) used for ChIP experiments was unable to recognize any amount of synthetic 18-mer CTD peptide with double phosphorylation site pT4pS5, whereas it could react with singly-phosphorylated CTD pT4 peptide in a linear dose-response fashion (**Fig. 1B**).

Next, we need to validate the specificity of phosphatase in order to establish that masking effect can be specifically removed. To do that, we cloned and purified *Drosophila* Ssu72/symplekin complex, and then treated doubly-phosphorylated pT4pS5 peptide. Dot blot showed that recognition of pT4 epitope was significantly reinforced by Ssu72/symplekin treatment, and sample treated with inactive enzyme showed no signal (**Fig. 1C**, **Suppl. Fig. S1A**). Ssu72 structure analysis highlights conformational features that enable high specificity of Ssu72 against pSer5 (**Suppl. Fig. S1B**), where the isomeric state of proline residues specified that only pSer5 fits in the active site, preventing the dephosphorylation of pT4 (*15, 16*). In addition, using a different approach, matrix-assisted laser desorption ionization-time-of-flight mass spectrometry (MALDI-TOF MS), we validated that Ssu72 acts selectively and effectively on removing Ser5 phosphosites, from both synthetic doubly-phosphorylated 18-mer peptide (pT4pS5) and singly-phosphorylated pS5 peptide (**Fig. 1D, Suppl. Fig. S1C**). At the same time, singly-phosphorylated pY1, pS2, pT4, and pS7 peptides were not affected in the result of the treatment at the concentration of phosphatase we used in the study (**Fig. 1D, Suppl. Fig. S1C**). Importantly, these results were reciprocated in *in vitro* experiment with whole cell lysates (HEK293), treated with active Ssu72 or catalytically dead mutant (**Fig. 1E, Suppl. Fig. S1D**). A time-course experiment showed that the enzyme starts to unmask pT4 as early as after 15-min of treatment, while signal was not changed in untreated cell lysate or the one treated by catalytically dead phosphatase (**Suppl. Fig. S1D***, right*).

Having validated Ssu72 activity and its ability to unmask pT4 sites for recognition, we leveraged this protocol to perform three biological replicates of ChIP-Seq experiments using HEK293 cell lysate separated into paired Ssu72 and mutant (Mut) samples and treated in a respective manner (**Fig. 1F**). As quality-control ‘fingerprint’ plots show, ChIP signal can be separated from the background signal (Input) in all three replicates, e.g. there was sufficient DNA enrichment due to antibody treatment (**Suppl. Fig. S1E**) (*17*). However, the shape of fingerprint curve is suggestive of broader domains instead of narrow peaks. Hence, we proceeded with calling peaks (--broad, *q* < 0.01) using MACS2 tool and identified whole-genome locations of pT4 peaks in Ssu72 and corresponding mutant samples (**Suppl. Fig. S1F**). Replicates were variable in numbers of called peaks, consistent with varying values of FRiP (fraction of reads in peaks) QC metric, however, 1% threshold recommended by ENCODE was met in every sample (*18*). Overall, metagene profiles (all hg19 genes) demonstrate increase in pT4 occupancy upon Ssu72 treatment, detected in all biological replicates (**Fig. 1F, Suppl. Fig. S1G**). Notably, unmasking in promoters and gene body was most consistent between biological replicates. As another ChIP quality control, we checked the distribution of called pT4 peaks over chromosomes (n of peaks normalized by chromosome length) in both conditions - Ssu72 and mutant. Resulting bar plots (**Fig. 1G, Suppl. Fig. 1H**) highlight unbiased ChIP, where signal (especially enriched upon Ssu72 treatment) was recorded on every chromosome. The highest normalized numbers of peaks were found on chromosome 19, which is not surprising taken into account its highest gene density - more than double the genome-wide average (*19*).

Differential binding (DiffBind) analysis identified 38,343 consensus affinity-based binding sites, e.g. pT4 sites merged between all replicates and conditions. Out of these consensus sites, 4,370 peaks were called as gained, and only 69 were depleted upon Ssu72 treatment (**Fig. 1H, Suppl. Table S1**). We used paired design with batch correction for DiffBind analysis and looser threshold for calling differential peaks (log_2_FC > 0.9, *p* < 0.05) due to the ‘broad’ nature of ChIP-signal. Principal component analysis (PCA) confirmed that differentially-bound peaks could discriminate samples into two conditions regardless of their batch, with the first PC accounting for 71% of variance (**Fig. 1I**). Hence, we could reveal 4,370 combinatorial pT4pS5 peaks in total, e.g. 11% of consensus pT4 peaks represent doubly-phosphorylated sites.

### pT4pS5 signal is predominantly contained within protein-coding genes

Having identified the genomic locations of pT4pS5 phosphosites, we asked what gene classes were mostly enriched with these marks. Profiling of pT4 signal after Ssu72 treatment separately on protein-coding (n = 19,838) and non-coding gene subsets (n = 40,426) revealed markedly low signal on non-coding genes, consistent between two top ChIP-enriched replicates (**Fig. 2A**, *left*, **Suppl. Fig. S2A**, *left*). The same difference was observed for paired mutant samples (**Fig. 2A**, *right*, **Suppl. Fig. S2A**, *right*). As we and others have previously reported, pT4 metagene profiles look different in protein-coding vs non-coding genes suggesting gene-specific role of pT4 (*13, 20*), and Ssu72 treatment was able to unmask some amount of pT4 signal in non-coding genes as well (**Suppl. Fig. S2B**). Yet, overall basal level of pT4 phosphorylation in non-coding genes was low even after Ssu72 treatment: in two replicates, TSS peak values hardly reached value of 0.05 (log_2_ read counts over Input), whereas FRiP-inferior replicate 3 showed only negative signal indistinguishable from background noise (**Suppl. Fig. S2B**). Interestingly, gene type-specific shape of metagene profile was not noticed when we re-analyzed published ChIP-Seq data on other CTD modifications, e.g. pSer5 and pSer2 (*21*) (**Suppl. Fig. S2C-E**).

**Figure 2.**
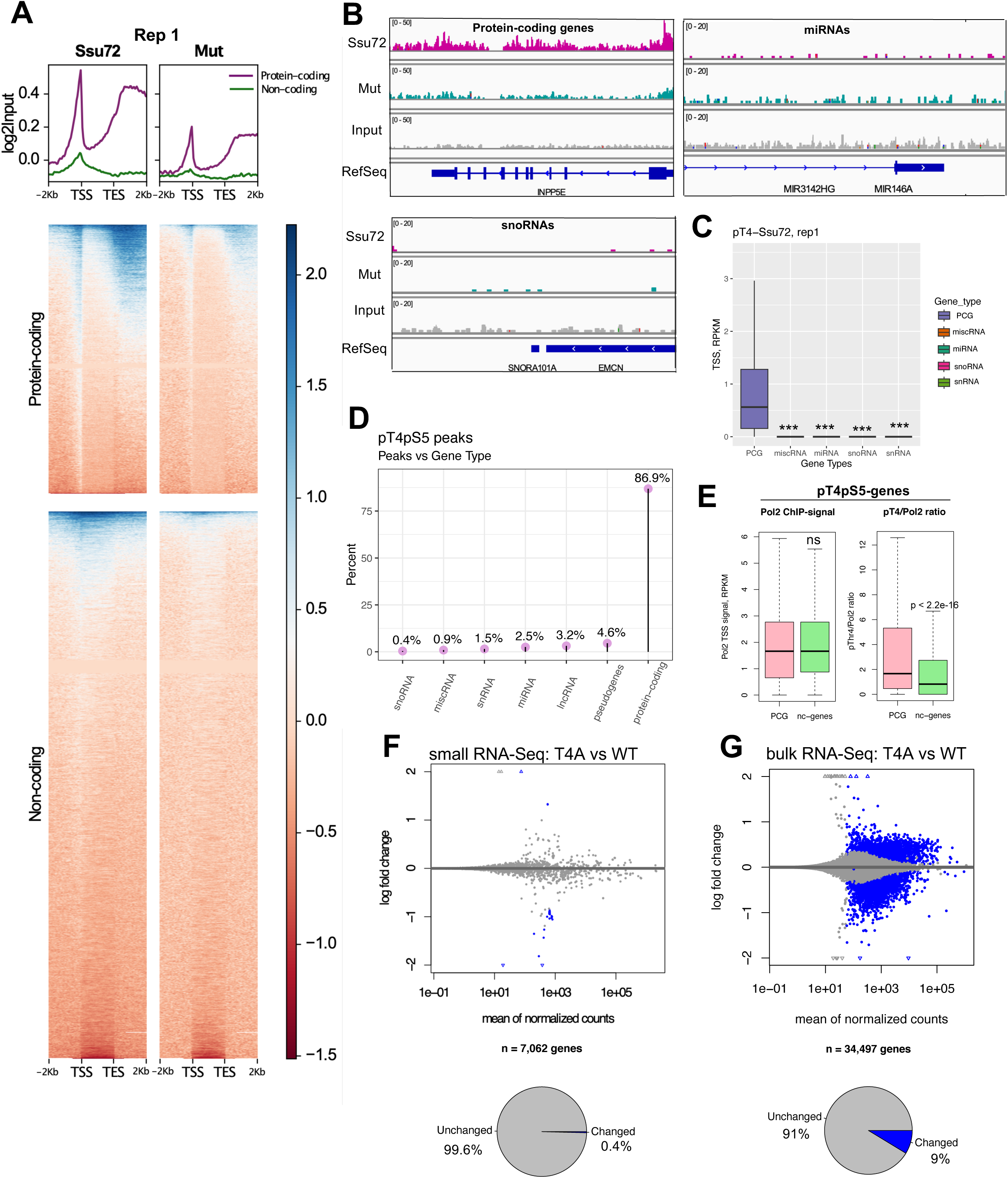
pT4pS5 double phosphorylation is contained within protein-coding genes in human cells. **A,** Whole-genome metagene profile and heatmaps of Input-normalized pT4 ChIP-Seq signal under Ssu72 vs Mutant treatment on protein-coding (n = 19,838) and non-coding subsets (n = 40,426) separately. Region between TSS and TES is scaled to 2,000 bp. Every bin is 50 bp. Heatmaps are sorted by mean value of each row (individual gene). One representative replicate is shown. **B,** Representative pT4 coverage IGV plots for a protein-coding gene (*INPP5E*), miRNA gene (*MIR146A*), and snoRNA gene (*SNORA101A*). **C,** TSS signal (RPKM) of pT4 signal upon Ssu72 treatment in 5 gene subsets: protein-coding (PCG, n = 19,838), miscRNAs (n = 2,034), miRNAs (n = 3,055), snoRNAs (n = 1,457), snRNAs (n = 1,916) derived from human genome annotation (gencode, hg19). Statistical comparison was performed using Wilcoxon test, \*\*\**p* < 2.2e-16. One representative replicate is shown. **D,** Lollipop chart of pT4pS5 peaks distribution across different gene types. **E,** Comparison of Pol2 ChIP-Seq signal (*left*) and normalized by Pol2 pT4 signal (representative replicate, *right*) on protein-coding (PCG, n = 2,471) vs non-coding (nc, n = 934) genes with pT4pS5 peaks. Comparison was performed using Wilcoxon test, \*\*\**p* < 2.2e-16, ns - not significant. **F-G,** Small (**F**) and bulk (**G**) RNA-Seq data visualization using MA plot (T4A vs wild-type cells). Pie charts showing relative number of deregulated genes (FDR < 0.05, both up- and down-regulated) in T4A vs WT is shown underneath.

The difference of pT4pS5 double phosphorylation in protein-coding and non-coding genes is obvious in the genomic mapping (**Fig. 2B**). Two exemplary genes from non-coding RNA category - *SNORA101A* and *MIR146A* - are shown in representative IGV plots in comparison with protein-coding gene (**Fig. 2B**). While the non-coding genes’ signal is barely above the noise with no difference with/without phosphatase treatment, protein-coding genes exhibit much higher enrichment of pT4 signal upon Ssu72 treatment. Deeper analysis of two superior ChIP-Seq replicates (1 and 2) showed that extracted RPKM values of pT4 ChIP-signal at genes’ starts were significantly lower (*p* < 2.2e-16) in all major classes of non-coding RNAs, including miscRNA, miRNA, snoRNA, and snRNA, versus PCG - protein-coding genes (**Fig. 2C, Suppl. Fig. S2F**). The annotation of identified pT4pS5 peaks (n = 4,370) pinpoints them almost exclusively (87%) within protein-coding genes, with little to no signal in non-coding genes (**Fig. 2D**). The same holds true for the high-confidence peak set containing filtered peaks with high enrichment in Ssu72-treated samples (log_2_FC > 1.5, n = 364 peaks) (**Suppl. Fig. S2G**).

To rule out the possibility that low pT4 signal in non-coding genes is a consequence of decreased Pol II recruitment to their promoters, we performed ChIP-Seq using RPB1-Pol II antibody to capture overall Pol II signal. Then, we normalized pT4 ChIP signal by overall RPB1-Pol II signal via calculating TSS ratios (RPKM pT4-Ssu72 / RPKM RPB1-Pol II) for every gene with detected combinatorial peaks (protein-coding, n = 2,471 genes vs non-coding, n = 934 genes) and plotted their distribution (**Fig. S2E, Suppl. Fig. S2H**). In the result, we did not observe decreased Pol II recruitment to the promoters of non-coding genes (median Pol II signal was 1.6610 in both subsets), and the normalized ratios were still significantly lower in non-coding genes (*p* < 2.2e-16), pointing out the true underlying difference in pT4pS5 occupancy. In addition, we filtered ultra-short protein-coding genes (< 1,000 bp, n = 451) and detected higher average pT4 signal over them in comparison with that of all sno and miRNA subset (n = 2,820) (**Suppl. Fig. S2I**). This analysis showed that very short protein-coding genes still have higher pT4 signal than non-coding ones.

Admittedly, even low signal-to-noise genomic signal can still lead to transcriptomic changes. To evaluate the importance of pT4 signal in expression of non-coding and protein-coding RNAs, we generated a synthetic construct of mammalian RPB1 with 52xCTD in which pT4pS5 marks cannot be generated. Since Ser5 mutation to alanine is known to result in cell lethality (*22*), and Ser5 residue is considered “unperturbable” in CTD study field, we use a construct where T4 in every heptad was replaced by alanine (T4A) in order to perturb pT4pS5 formation.

We subjected transfected mutant cells to 72-h α-amanitin treatment to shut-off endogenous RNA Pol II and measured non-coding and protein-coding RNA expression via small RNA-Seq (sRNA-Seq) and bulk RNA-Seq with polyA enrichment, respectively. Interestingly, very few non-coding RNAs (only 28 out of 7,062 annotated genes) got deregulated (FDR < 0.05) upon Thr4 perturbation vs wild-type (**Fig. 2F, Suppl Fig. S2J**). Most deregulated (decreased) small RNAs belonged to a lesser studied class of non-coding RNAs called Y-RNAs, among which 31 pT4pS5 marks were identified in their genome sequence. As a comparison, bulk RNA-Seq estimated 9% of expressed genes (2,998 out of 34,497 with non-zero counts) to be deregulated (FDR < 0.05) upon T4A mutation (**Fig. 2G**), the majority of which were indeed annotated as protein-coding. RNA-Seq library could also capture 8,724 polyA-containing lncRNAs which is about ∼50% of annotated lncRNAs in human genome (hg38: n = 16,876). Similar to small RNAs, only minor percent of lncRNA (n = 69, 0.8%) changed expression upon T4 perturbation (**Suppl Fig. S2K)**.

Taken together, CTD’s pT4pS5 occur during eukaryotic transcription as a protein-coding gene specific modification on RNA polymerase II.

### pT4pS5 combinatorial marks are enriched on neurogenesis-related genes

Now that we established that pT4pS5 only occurs during protein-coding genes’ transcription, we sought to identify its biological function. Based on the whole-genome profiling of pT4 upon Ssu72/Mutant treatment, we noticed most consistent enrichment in gene regions corresponding to promoters and gene body. To assess statistical significance of this enrichment, we calculated three metrics - TSS signal, mean intragenic (e.g., gene body) signal, and TES signal - for every protein-coding gene, and then compared resulting values in a pairwise fashion between Ssu72 and catalytically-dead mutant (**Fig. 3A, Suppl. Fig. S3A**). The difference between conditions was highly significant (*p* < 2.2e-16) in every biological replicate for promoters and gene body, but not TES region. Exemplary genes with pT4pS5 double phosphorylation in promoters and intragenic regions, respectively, are shown as IGV plots beneath (**Fig. 3A***, lower*).

**Figure 3.**
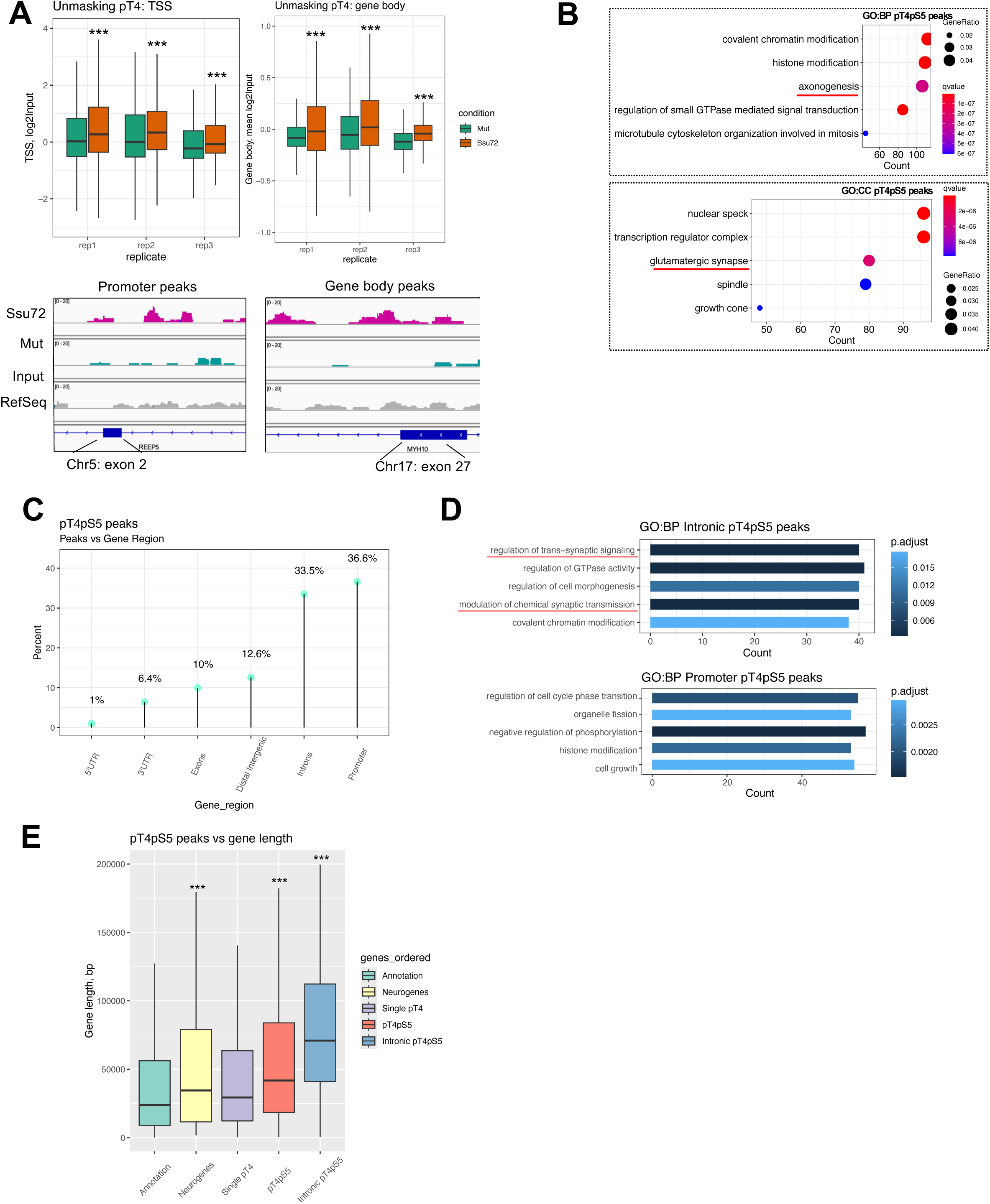
pT4pS5 signal is enriched on protein-coding genes related to neurogenesis. **A,** TSS (*left*) and gene body (*right*) Input-normalized pT4 signal (protein-coding genes, n = 19,838) upon treatment with either Ssu72 or Mutant in all performed replicates. Statistical comparison was performed using Wilcoxon test, \*\*\**p* < 2.2e-16. Exemplary IGV plots of promoter and gene body peaks are shown underneath. **B,** Gene ontology (GO) analysis of biological processes (BP) and cellular components (CC) that were enriched among genes with pT4pS5 peaks. Analysis was performed using ‘clusterProfiler’ package in R and all human genes as gene universe, with BH-corrected FDR. Neuron-related terms are highlighted in red. **C,** Lollipop chart of pT4pS5 peaks distribution across different gene regions. **D,** GO analysis of biological processes that were enriched among genes with pT4pS5 peaks, located in introns (*upper*) or promoters (*lower*). Analysis was performed using ‘clusterProfiler’ package in R and all human genes as gene universe, with BH-corrected FDR. Neuron-related terms are highlighted in red. **E,** Genes’ lengths in 5 gene subsets - annotated human protein-coding genes (“Annotation”, n = 19,966), annotated neurogenes from GO:0050769 (“Neurogenes”, n = 475), genes with single pT4 peaks (n = 10,648) and double pT4pS5 peaks (n = 2,471), and intronic pT4pS5 (n = 832) separately. Comparison vs Annotation was performed using Wilcoxon test, \*\*\**p* < 2.2e-16. Only protein-coding genes were analyzed and plotted.

To delineate the biological processes for which pT4pS5 marks may be important, we proceeded with functional annotation of genes with identified Ssu72-enriched peaks (n = 4,370) (**Fig. 3B**). Gene ontology analysis identified chromatin modification and axonogenesis as top-3 enriched biological processes, and nuclear speck (or splicing speck), transcription regulator complex, and synapse as top-3 enriched cellular components, FDR < 0.05 (**Fig. 3B, Suppl. Table S2**). These results led us to hypothesize that pT4pS5 pattern may be important for transcription of neurogenesis-related genes. In support of this hypothesis, neurogenesis-related biological processes were significantly enriched among genes with pT4pS5 peaks called as differentially-bound using different combinations of ChIP-Seq biological replicates, e.g. replicates 1 and 2, replicates 2 and 3, etc. (**Suppl. Fig. S3B**). In addition, the high-confidence pT4pS5 peak set (log2FC > 1.5) also pinpointed GO:0050769 ‘positive regulation of neurogenesis’ among its top-3 upregulated processes (**Suppl. Fig. S3C**).

To gain more insight from pT4pS5 ChIP signal, we annotated Ssu72-enriched peaks based on the gene region (**Fig. 3C**). The majority of peaks belonged to promoter regions, as expected, considering that ‘masking’ adjacent pSer5 has a big peak at genes’ starts. The second most prevalent category was intronic pT4pS5 peaks (**Fig. 3C, Suppl. Fig. S3D**). We do not observe high percent of pT4pS5 peaks in 3’-UTRs, probably due to absence of pSer5 in 3’-ends of the genes. For comparison, about 6% of single pT4 peaks were located in 3’-UTRs (*13*).

Intriguingly, when we compared functional enrichment among intronic-mapped and promoter-mapped pT4pS5 peaks, we noticed neurogenesis-related biological processes only among intronic peaks, suggesting there may be downstream neurogenesis regulators being recruited directly or indirectly to intronic pT4pS5 phosphosites (**Fig. 3D**). Having noted ‘covalent chromatin modification’ as another biological process assigned to intronic pT4pS5 peaks, we calculated %overlaps of these peaks and activating histone marks from large-scale Roadmap Epigenomic project (*23*), with neuronal progenitor cells as the query (**Suppl. Fig. S3E**). Interestingly, >75% of intronic pT4pS5 peaks overlap with activating histone marks, suggesting that these genomic regions are important transcriptional ‘hot spots’.

It has been noted that neuronal genes are especially long (*24*), and longer genes are more likely to have brain-related functions and participate in synaptic pathways (*25*). Considering these facts in conjunction with enrichment of synapse-related functional terms among genes with intronic pT4pS5 peaks, we analyzed lengths of genes with detected single and double phosphosites (**Fig. 3E**). Importantly, genes with pT4pS5 peaks, especially intronic-mapped, were much longer in comparison with annotation (*p* < 2.2e-16), whereas genes with single pT4 peaks (not enriched upon Ssu72 treatment, n = 33,845 peaks, 10,648 protein-coding genes) were not markedly longer than annotated subset of all protein-coding genes (**Fig. 3E**). Consistently, neuronal genes from GO:0050769 group (‘positive regulation of neurogenesis’) were also longer (*p* = 1.1e-14) than annotation metric (**Fig. 3E**). Thus, pT4pS5 marks are enriched in the intronic region of longer genes, frequently the ones involved in neurogenesis.

Taken together, all this evidence highlights the functional impact of pT4pS5 phosphosites on the transcriptional activity of neurogenesis-related genes.

### Transcription Export (TREX) complex components are recruited via pT4pS5 combinatorial marks

To test our hypothesis that neurogenesis mediator(s) can be recruited to pT4pS5 CTD double phosphosites, we designed an *in vitro* label-free pulldown experiment. Although a few kinases have been nominated as a putative Thr4 kinase, such as P-TEFb and PLK3 (*14*), there is no known kinase that can phosphorylate the Thr4 position in human cells. Thus, to circumvent this barrier we purified a 26X mutant CTD construct where every Thr4 was mutated to glutamate (T4E) to mimic the negative charge that would be present if the Thr4 residue was phosphorylated at this position. The construct also contains an N-terminal hexahistidine glutathione S-transferase (HGST) tag, which can bind to glutathione beads to immobilize our CTD “bait” in the pulldown experiment. Using the mutant T4E 26X yeast CTD construct and a 26X wildtype yeast construct as a control, we treated the substrates with an Erk2 kinase, which only phosphorylates Ser5 *in vitro* based on our previous mass spectrometry data (*26, 27*). Preliminary MALDI-TOF mass spectrometry analysis showed that Erk2 treatment was able to add 14-29 phosphate groups to wild-type 26xGST CTD peptide and 13-28 phosphate groups to mutant 26xGST-T4E CTD peptide, depending on respective biological duplicate (**Fig. 4A, Suppl. Fig. S4A-E**). The addition of phosphate groups to the mutant T4E CTD by a kinase with Ser5 specificity suggests that we were able to generate the T4EpSer5 combinatorial mark *in vitro*.

**Figure 4.**
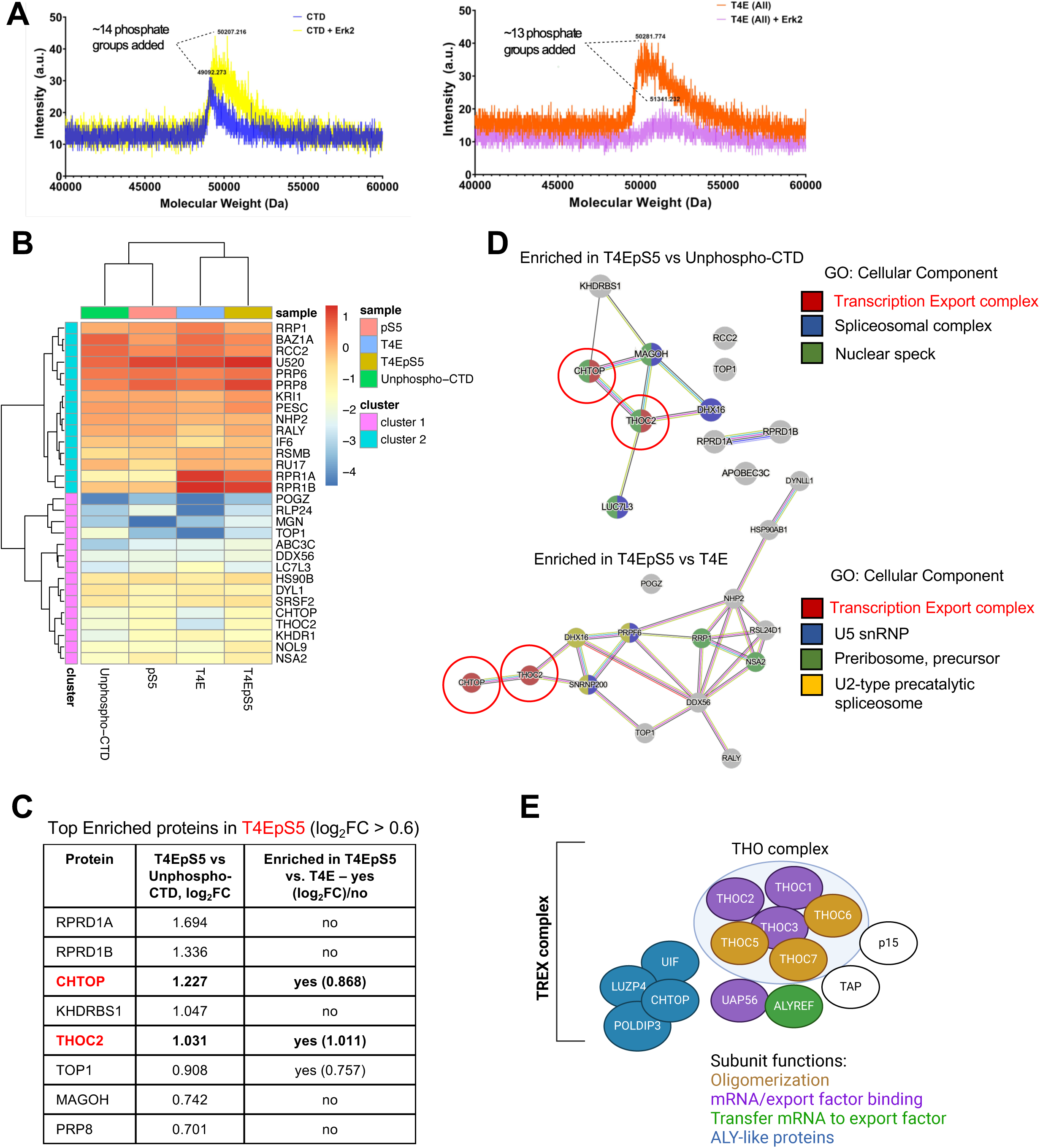
TREX complex is associated with pT4pS5 double marks. **A,** MALDI-TOF profiles of wild-type 26xCTD construct (*left*) and T4E-mutated 26xCTD construct (*right*) before and after phosphorylation reaction using Erk2 kinase (2.4 µM). One representative replicate is shown. **B**, A heatmap of normalized protein spectral counts in 3 conditions: pS5, T4E, T4EpS5 vs unphospho-CTD (replicate 1). Only proteins that were enriched (cut-off log_2_FC > 0.6) in pulldown at least in one group (pS5, T4E, T4EpS5) versus unphosphorylated control are shown. Statistical comparison was performed using ‘limma’, and hierarchical clustering was performed using ‘pheatmap’ in R. **C**, A table with top-enriched proteins in T4EpS5 samples vs unphosphorylated control (cut-off log_2_FC > 0.6). **D**, PPI (protein-protein interactions) networks of proteins enriched in T4EpS5 pulldown vs unphosphorylated control (*upper*) or vs single T4E (*lower*). Networks were built using the STRING database; GO analysis (CC: cellular component) was performed using built-in ‘Analysis’ tab. **E**, Molecular composition of TREX complex (*30, 48*).

For the ‘bait’ in our pulldown experiment, we treated the wildtype and mutant T4E 26X CTDs with Erk2 kinase and had both the untreated wildtype and mutant T4E CTDs as controls. The treated and untreated CTD constructs were incubated with the ‘prey’ HEK293 nuclear lysate. The samples were subjected to a series of washes to reduce non-specific binding and eluted in preparation for MS analysis. To focus on proteins that were specifically enriched in T4EpSer5 interactome, the abundances of hit proteins in T4EpSer5 samples were compared to that in all other conditions: unphosphorylated control, T4E, and pSer5. Briefly, we applied median centering on clean and decontaminated log2-transformed PSM (peptide spectral match) counts for every experimental sample (8 samples, 4 conditions in total) in order to normalize the replicates, and then we fitted linear models with different contrasts using limma in R (*28, 29*). The normalized and filtered dataset can be found in **Suppl. Table S3**.

To identify proteins enriched in T4EpSer5 samples, we selected the proteins with at least 1.5-fold (log_2_FC > 0.6) enrichment in T4EpSer5 vs. other conditions (T4E, pSer5, Control) and merged them into a new dataset with unique rows corresponding to enriched proteins (**Fig. 4B**). Hierarchical clustering separated proteins into higher- and lower-expressed groups; sample-wise, T4E and T4EpSer5 samples were clustered together (**Fig. 4B**). Interestingly, when we ranked proteins enriched (log_2_FC > 0.6) in T4EpSer5 samples compared to the unphosphorylated control, we noticed two proteins - CHTOP and THOC2 - from the multi-protein complex TREX (TRanscription and EXport) (*30*) amongst the top-5 ranks (**Fig. 4C**). THOC2 and CHTOP were also the two most enriched in T4EpSer5 pulldown compared to singly-modified T4E. The presence of RPRD1A and RPRD1B amongst top-enriched proteins associated with T4E, but not specifically with T4EpS5, also provides further validation of our proteomic studies since our previous biophysical characterization reveals that RPRD1A/B bind to pT4 tightly but the association disrupted upon flanking pSer5 (*13*).

To gain insight into the functional relations between identified proteins, we built several interaction networks based on protein-protein interactions (PPI) using the STRING database (https://string-db.org/). We used protein lists enriched in T4EpSer5 versus 1) unphosphorylated control; 2) T4E; 3) pSer5 (**Fig. 4D, Suppl. Fig. S4F**). Next, we applied GO: CC (Gene Ontology: Cellular Component) analysis to the networks: as a result, TREX complex was highlighted in 2 out of 3 protein networks (**Fig. 4D, Suppl. Fig. S4F**). Interestingly, spliceosome and nuclear speck cellular components (MAGOH, PRP8) were also enriched in T4EpSer5 pulldown versus unphosphorylated control, yet they were not different between single T4E and T4EpS5 (**Fig. 4C**). These facts align with the notion that multi-protein TREX complex (its composition is shown in **Fig. 4E**) is important for mRNA splicing and proper neurodevelopment (*31–33*). In order to additionally validate the idea of TREX recruitment to CTD of Pol II, we expressed exogenous GFP-tagged Pol II in U2OS cells, and then performed immunostaining using nuclear stain (DAPI), GFP-antibody, and THOC2 antibody (**Suppl. Fig. S4G-I**). We observed high amount of co-localization between Pol II and THOC2, a TREX component, spatiotemporally organized in the form of nuclear speckles.

Overall, our proteomic analysis has uncovered a potential proteome-level link between doubly-phosphorylated CTD pattern and TREX complex.

### pT4pS5 controls expression and nuclear export of long transcripts of neurogenesis-related genes

To characterize functional interactions between TREX complex and pT4pS5 CTD pattern, we first turned to available external omics data on TREX components, starting with CHTOP and THOC2. Whole-genome profiling of Flag-CHTOP ChIP-Seq binding shows that CHTOP is recruited to gene promoters (GSE130992) (**Fig. 5A**, *left*). Annotation of CHTOP-peaks shows that 39%-48% of peaks map to intronic regions (**Fig. 5A**, *right*, **Suppl. Fig. S5A**). Consistent with published importance of TREX complex in proper neurodevelopment (*33*), functional annotation (GO:CC) of both biological replicates of CHTOP peaks highlights synapse-related terms (**Fig. 5B, Suppl. Fig. S5B**), with ‘glutamatergic synapse’ reminiscent of top-cellular component genes having double pT4pS5 peaks (**Fig. 3B**). Most frequent introns that have CHTOP peaks are 1-3 introns consistent with co-transcriptional mRNA processing function of TREX (**Fig. 5C**, *left*, **Suppl. Fig. S5C**). Intriguingly, a subset of pT4pS5 intronic peaks also follows the same distribution (**Fig. 5C**, *right*), overlapping with CHTOP peaks at some genomic loci (**Fig. 5D**).

**Figure 5.**
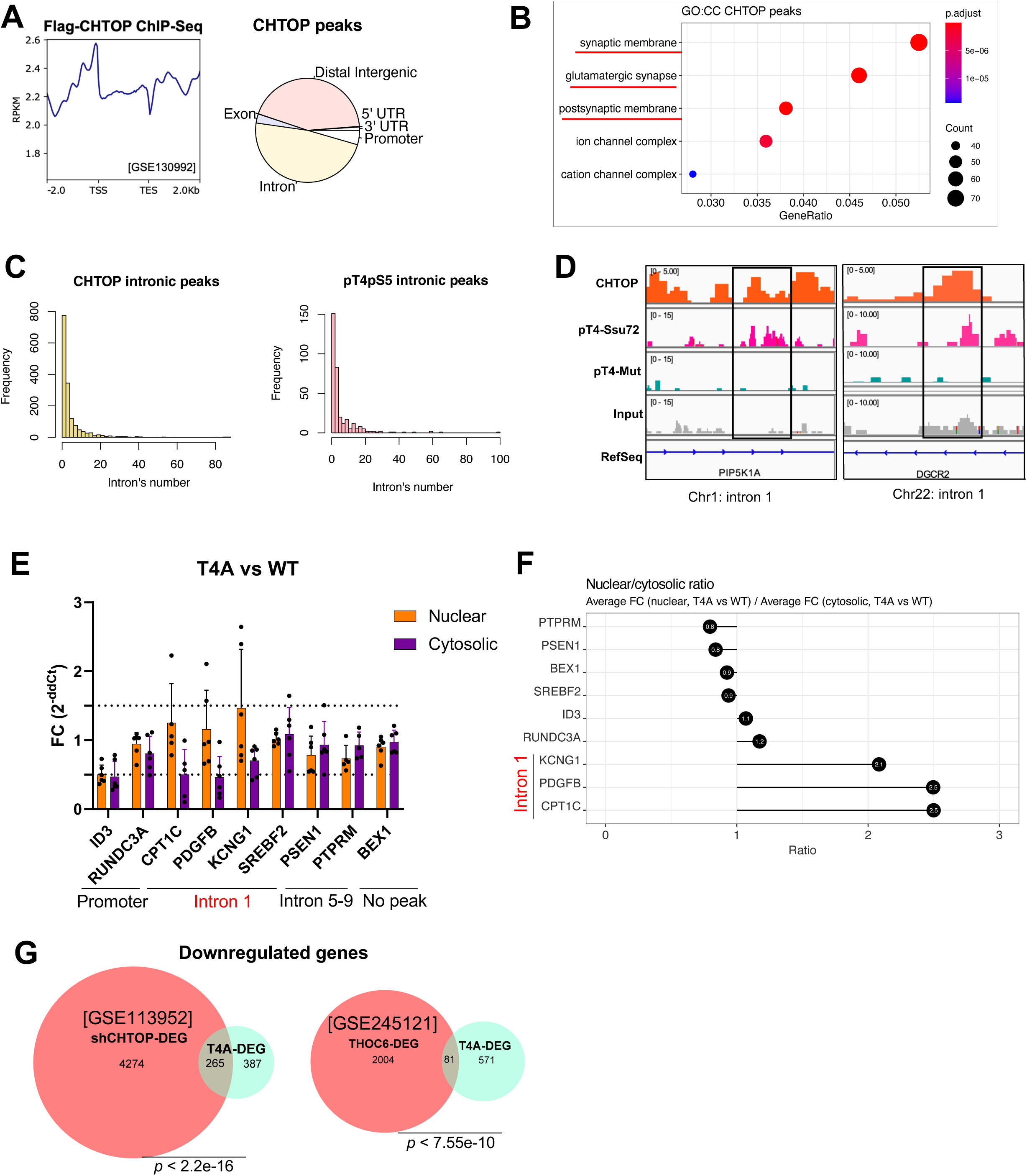
Intronic-mapped pT4pS5 peaks recruit TREX for processing and nucleocytoplasmic export of mRNAs. **A,** Metagene profile of Flag-CHTOP ChIP-Seq signal on all protein-coding genes (n = 19,984) [GSE130992] (*left*) and annotation of MACS3-called (*p* < 0.001) CHTOP peaks (*right*). Region between TSS and TES is scaled to 2,000 bp. Every bin is 50 bp. Peak annotation was performed using ‘chipseeker’ R package. One representative replicate of peaks is shown. **B**, GO analysis of cellular components enriched among genes with CHTOP peaks [GSE130992]. Analysis was performed using ‘clusterProfiler’ package in R and all human genes as gene universe, with BH-corrected FDR. One representative replicate of peaks is shown. **C**, Frequency histograms of CHTOP [GSE130992] (*left*) and pT4pS5 (*right*) intronic peaks broke down by intron’s number. One representative replicate of CHTOP peaks is shown. **D**, Colocalization of CHTOP and pT4pS5 peaks at select genomic loci visualized by IGV. **E**, RT-qPCR measurements of nuclear and cytoplasmic levels of select transcripts with (*ID3, RUNDC3A, CPT1C, PDGFB, KCNG1, SREBF2, PSEN1, PTPRM*) or without (*BEX1*) pT4pS5 peaks. Shown are mean with SD derived from 5-6 biological replicates. Analysis (T4A vs wild-type) was done using ddCt method normalized by *GAPDH* gene expression. **F**, Nuclear/cytosolic ratios defined as the ratios between average nuclear and cytosolic Fold Changes (T4A vs wild-type) in transcript abundance across tested gene panel. Ratios were ranked in ascending order. Enrichment of genes with intron1-mapped pT4pS5 peaks among those with impaired nucleocytoplasmic export is highlighted. **G**, Overlap analysis of downregulated genes (FDR < 0.05, log_2_FC < -0.6) upon T4A mutation and CHTOP-knockdown [GSE113952] (*left*) and THOC6-LOF [GSE245121] (*right*). Statistical significance of overlaps was tested using Fisher’s exact test for count data.

It has been known that TREX plays a major role in the functional coupling of different steps during mRNA biogenesis and nuclear export, as well as cell differentiation and development, likely because this complex regulates the expression of key developmental genes (*34*). Both THOC2 and CHTOP genes are highly important for neurogenesis, as ClinVar database (*35*) includes several pathogenic, likely pathogenic, and VUS (variants of unknown significance) germline mutations in *CHTOP* and *THOC2* genes associated with X-linked intellectual disability (**Suppl. Fig. S5D**). Therefore, assuming the interaction of pT4pS5 marks with TREX component(s), we hypothesized that loss of pT4pS5 in cells would also cause impairment of mRNA export out of the nucleus. To test this hypothesis, we collected T4A and wild-type cells after 72-h α-amanitin treatment, extracted total RNA from nuclear and cytosolic cellular fractions separately, and tested the amount of select transcripts using qPCR (**Fig. 5E**). As a “purity control”, we performed Western blotting of histone H3, which showed up preferentially in the nuclear fraction as expected (**Suppl. Fig. S5E**). In our gene panel, we included those genes that had promoter pT4pS5 peaks (*ID3, RUNDC3A*), intron-mapped pT4pS5 peaks located either in intron 1 that was the most preferential location (*CPT1C, PDGFB, KCNG1, SREBF2)* or in intragenic introns (*PTPRM* - intron 5*, PSEN1* - intron 9). Many of these genes are expressed in discrete brain areas and required for neurogenesis and/or synaptic plasticity (*36–39*). We also included a gene without detected double phosphorylation (*BEX1*) as negative control (**Fig. 5E, Suppl. Fig. S5F**). We normalized the relative expression by *GAPDH* level, which did not contain pT4pS5 mark and whose expression was stable across fractions and conditions (**Suppl. Fig. S5G**). Our results show that nucleocytoplasmic export of transcripts with pT4pS5 mark in the first intron is hindered in T4A cells compared to control: their transcripts tended to accumulate in the nucleus, yet show reduction in the cytosolic fraction, with average nuclear/cytosolic ratios of Fold Changes over control > 2 (**Fig. 5E-F**). At the same time, nucleocytoplasmic export of other transcripts was largely unaffected.

We next asked if perturbation of T4 phosphorylation also reduces overall expression of specific mRNA transcripts that are functionally controlled by TREX. To answer this question, we overlapped markedly downregulated genes (FDR < 0.05, log_2_FC < -0.58, **Suppl. Table S4**) upon T4A mutation and downregulated genes from two other recent studies - in CHTOP-knockdown HEK293 cells (GSE113952) and THOC6 loss-of-function neuroprogenitor cells (GSE245121) (*33, 40*). In both cases, gene overlap was significant (*p* < 2.2e-16 and *p* < 7.55e-10, respectively) indicative of a tight correlation between pT4 and TREX transcriptional roles (**Fig. 5G**). TREX complex is necessary for RNA processing associated with corticogenesis which becomes impaired in THOC6 intellectual disability syndrome (*33*). Consistently, pooled downregulated DEGs with pT4pS5 peaks are enriched for axonogenesis-related genes (**Suppl. Fig. S5H**). Using qPCR, we experimentally validated downregulation of exemplary *POLA1* gene upon both T4 and THOC2 perturbation (average shTHOC2 knockdown efficiency ∼80% estimated by Western blotting) in HEK293 cells (**Suppl. Fig. S5I-J**). *POLA1* (DNA polymerase a-primase) plays a crucial role in DNA replication and is one of causative genes in X-linked syndrome involving intellectual disability (*41*).

Taken together, this data highlights that pT4pS5 marks recruit TREX complex to the ongoing transcription of certain genes enriched for those related to neurogenesis for their processing and nucleocytoplasmic export.

### Transcriptional outcome of pT4 and TREX perturbation show overlapping mis-splicing of longer genes related to neurogenesis

Another important function of TREX is co-transcriptional splicing, with a particularly significant role in the splicing of neuronal genes (*33*). We performed canonical alternative splicing (AS) analysis with rMATS (*42*) on RNA-Seq data from mutant HEK293 T4A cells vs wild-type cells (*13*). As a result, pT4 loss led to alteration in splicing site (**Suppl. Fig. S6A***, left*). Compounded with the potential role of pT4pS5 in recruiting TREX to ongoing transcription, we asked if combinatorial pT4pS5 marks play a role in mediating splicing through TREX. First, to assess the significance of association between pT4pS5 peaks and splicing, genes were separated into ‘properly-spliced’ and ‘mis-spliced’ in T4A condition vs wild-type (FDR < 0.05, ILD, inclusion level difference > 10%) based on AS analysis. Based on peaks, genes were separated into the categories with detected pT4 peaks and without them (**Fig. 6A**). The resulting contingency table was visualized as mosaic plot, colored by Pearson residuals of chi-squared test (X-squared = 1739.7, overall *p* < 2.2e-16) (**Fig. 6A**). As the plot shows, genes with pT4 phosphosites were significantly overrepresented among mis-spliced genes upon T4A mutation, e.g. pT4 signal is significantly associated with proper canonical splicing. In addition, we confirmed direct interaction of pT4 with a known spliceosomal factor PRPF8 via co-immunoprecipitation experiments using 26X T4E constructs with or without treatment with Erk2 kinase as ‘bait’ and HEK293 nuclear lysate as ‘prey’ (**Suppl. Fig. S6B**). This is consistent with our pulldown results, where we detected PRPF8 as a protein interacting with both T4EpS5 and T4E (**Fig. 4C**).

**Figure 6.**
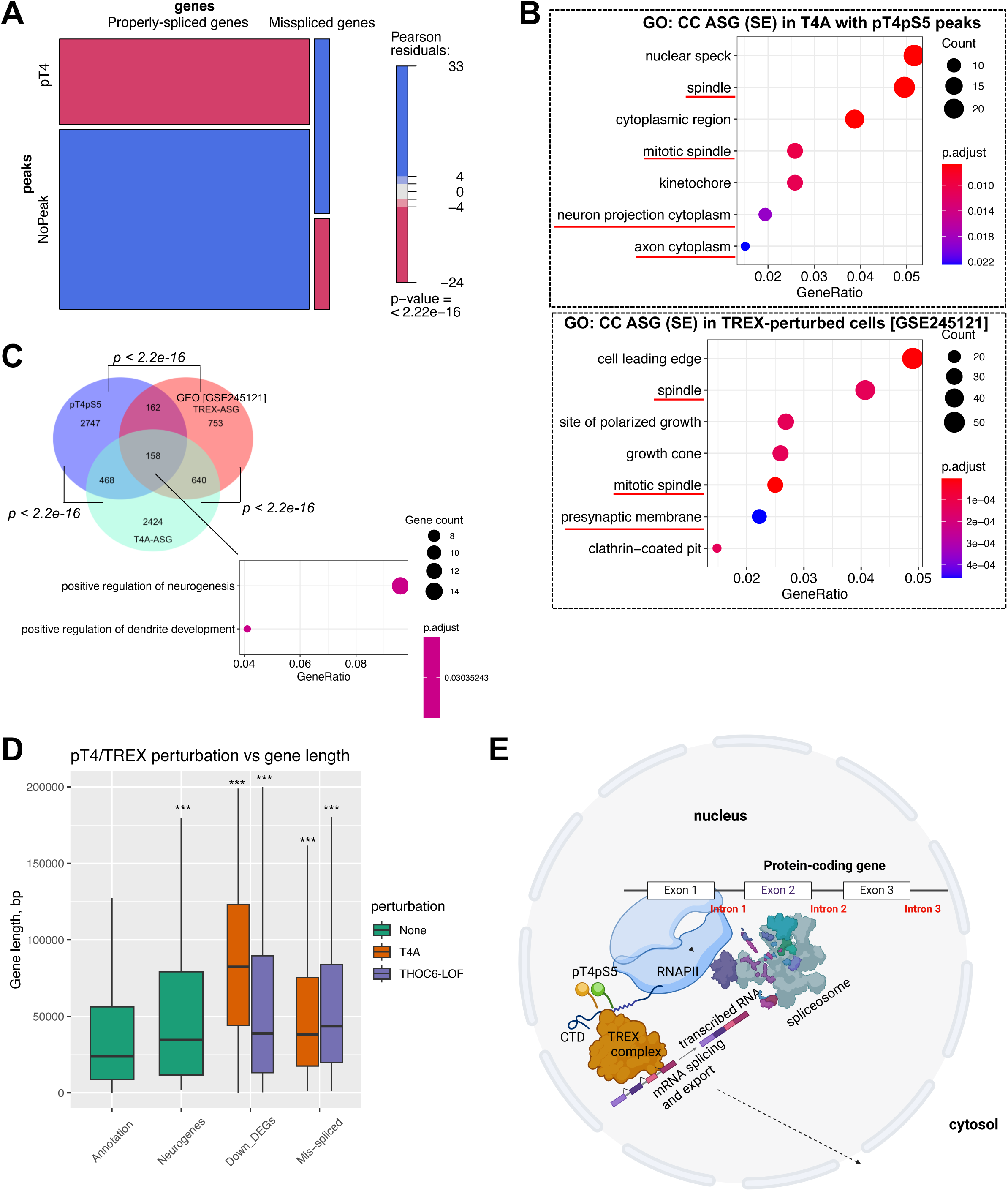
The pT4pS5 marks drive splicing of lengthy neurogenesis-related genes through TREX. **A,** Mosaic plot illustrating significant association (*p* < 2.2e-16) between two categorical variables - splicing (Properly-spliced vs Mis-spliced genes) and pT4 peaks (pT4 vs NoPeak). Statistical testing was performed using Chi-squared test. **B**, GO analysis of cellular components enriched among alternatively-spliced (SE, Skipped Exon category) genes upon T4 perturbation (*upper*) or TREX perturbation [GSE245121] (*lower*). Only genes with pT4pS5 double phosphorylation peaks were used for upper plot. Analysis was performed using ‘clusterProfiler’ package in R and all human genes as gene universe, with BH-corrected FDR. Neuron-related terms are highlighted in red. **C**, Overlap analysis of three gene subsets: genes with pT4pS5 peaks, alternatively-spliced (ASG) upon T4A and TREX perturbation (THOC6-LOF, GSE245121). Significance of overlaps was tested using Fisher’s exact test for count data. GO analysis (*below*) highlights neurogenesis as biological process uniquely associated with shared genes (n = 158). **D**, Lengths of downregulated (n = 627, n = 1,809) and mis-spliced (n = 3,694, n = 1,580) genes upon T4 or TREX perturbation, respectively, in comparison with annotated human protein-coding genes (“Annotation”, n = 19,966) and annotated neurogenes from GO:0050769 (“Neurogenes”, n = 475). Comparison vs Annotation was performed using Wilcoxon test, \*\*\**p* < 2.2e-16. Only protein-coding genes were analyzed and plotted. **E**, A model illustrating how intronic-mapped pT4pS5 combinatorial phosphorylation on CTD recruits TREX complex and thereby mediates co-transcriptional mRNA splicing and export of lengthy protein-coding mRNAs. The model was created using BioRender software.

Since inclusion or skipping of exons are much more prevalent compared to other types of alternative splicing events (∼10 fold) under both perturbations - T4A mutation and THOC6 loss-of-function (THOC6-LOF, GSE245121) (**Suppl. Fig. S6A**), we functionally annotated genes that underwent exon inclusion/skipping (‘SE’) upon T4A/TREX perturbation (**Fig. 6B**). Among the top enriched GO terms (CC: cellular component), we found cell cycle and neuronal gene categories (**Fig. 6B**). Importantly, genomic ranges of pT4pS5 peaks and SE events (‘upstream exon start’ - ‘downstream exon end’) directly overlapped by 35% (230/660 pT4pS5 ranges, lifted to hg38), indicating strong correlative effect of pT4pS5 signal on the choice of alternate exons by the splicing machinery at least in 1/3 cases.

When we overlapped three gene subsets (1 - genes with pT4pS5 peaks; 2 - genes with AS changes in THOC6-deficient human neuroprogenitor cells; 3 - genes with AS changes in T4A-mutated HEK293 cells), we identified significant gene overlap in all three statistical tests (Fisher’s exact test *p* < 2.2e-16, with 34,497 genes having non-zero counts in T4A/Control library as gene universe) (**Fig. 6C**). Noticeably, genes composing an overlap of all three datasets (n = 158) were functionally important for neurogenesis process (**Suppl. Table S5, Fig. 6C**). Among these genes, one-half (70/158) had pT4pS5 peaks mapped to introns, which was the most prevalent location of TREX and pT4pS5 peaks. Within overlapping mis-spliced genes (n = 798) between T4A and THOC6-LOF, 115 AS events were exactly identical in terms of both direction of event and its genomic coordinates (**Suppl. Table S6**). To further assess the colocalization between pT4pS5 CTD-pattern and TREX, we correlated binned ChIP-signal of CHTOP (GSE130992) and our pT4-Ssu72 dataset on two gene subsets - 1) all protein-coding genes and 2) genes composing the pT4pS5/T4A/THOC6 overlap (n = 158) from Fig. 6C (**Suppl. Fig. S6C**). In the result, correlation coefficients, although weak, became stronger (*p* = 7.99e-0.5) when we filtered the queried gene subset.

Analysis of recently published ChIP-Seq and TT-Seq datasets (*40, 43*) suggest TREX components are coupled to transcription and their perturbation affects transcription metrics, such as Pol II pausing and elongation speed (**Suppl. Fig. S6D**, *left*; **Suppl. Fig. S6E-G**). Importantly, perturbation of a subunit of TREX, THOC5 (GSE173374) and Thr4 phosphorylation (T4A, GSE37519) both lead to Pol II stalling on gene promoters (**Suppl. Fig. S6D**). Intrigued by this shared phenotype, we set out to explore which gene categories preferentially get deregulated, e.g. become differentially expressed (downregulated) or mis-spliced, upon either of these perturbations.

Thus, we accessed THOC6-LOF dataset (GSE245121) again and compared the deregulated genes’ lengths and T4A-deregulated genes’ lengths with annotation (all protein-coding genes, n = 19,966) metric (**Fig. 6D**). As the boxplot shows, upon both perturbations, we observe significantly longer genes being downregulated (FDR < 0.05, log_2_FC < -0.58) and/or alternatively spliced (FDR < 0.05, ILD > 10%) (*p* < 2.2e-16). Interestingly, neuronal genes from GO:0050769 group (‘positive regulation of neurogenesis’) were also generally longer (*p* = 1.1e-14) than annotation metric (**Fig. 6D**). In addition, upon THOC5 perturbation (GSE37519), longest genes had the most marked Pol II stalling on the promoters (**Suppl. Fig. S6G***, right*), indicative of TREX importance for active transcription of longer genes. This result echoes our observation that intronic-mapped pT4pS5 peaks preferably locate in the longest genes (**Fig. 3E**). Finally, using SpliceTools suite (*44*), we estimated that the sizes of skipped/included and downstream exons upon T4A mutation were slightly longer than annotation metric, but comparable with individual RBP knockdown datasets from SpliceTools collection (**Suppl. Fig. S6H**).

Taken together, this data indicates that pT4pS5 phosphosites, especially intronic-mapped, are significantly associated with proper splicing of long neurogenesis-related genes, probably through their association with TREX complex.

## DISCUSSION

Several *in vitro* biophysical studies revealed that transcription regulators can exhibit preferential binding to some combinations of PTM patterns on the CTD. For example, the double mark pS2/pS5 is directly recognized by the splicing factor U2AF65 (*45*). In another example, yeast termination factor Rtt103p binds pS2 and pT4 CTD well, but doubly-phosphorylated pS2/pT4 pattern diminishes its binding affinity (*46*). However, the physiological relevance of such preferential binding remains elusive, even though combinational PTM patterns have been detected in cells (*10, 11*). To identify the combinatorial PTMs of CTD in a whole-genomic fashion, we exploited a common phenomenon of interference of nearby phosphorylation with antibody recognition (e.g., antibody masking). We used a highly selective phosphatase to remove the masking effect and based on the comparison of signal with and without unmasking, we could deduce the location of the double phosphorylation patterns (**Fig. 1A**). We applied this framework to the detection of pT4pS5 double phosphosites.

Our study determined that the absolute majority of pT4pS5 marks are contained within protein-coding genes in human cells, most preferably in the gene promoters and introns. We observed few combinatorial pT4pS5 marks in non-coding genes, and little to no change in their expression levels upon T4 mutation. Importantly, we found that combinatorial pT4pS5 do not appear in downstream gene regions, potentially due to the prominent decline of pS5 in 3’-ends (*47*). In contrast, pT4 remains highly abundant after TES (*13*). Thus, for the first time, we mapped the whole-genomic location of double phosphorylation pattern on CTD and our method should be applicable to detect all the physiologically important combinatorial CTD marks.

The exclusive detection of pT4pS5 on protein-coding genes suggests such code as gene-specific, which is subsequently supported by our functional ‘late’ integration of omics data (genomic, proteomic & transcriptomic) and validation experiments. The pT4pS5 patterns over protein-coding genes are found to associate with TREX, a key regulator for co-transcriptional splicing and nucleocytoplasmic export of mRNAs, especially critical for neurogenesis-related biological processes. Notably, TREX subcomplex, THO, has been proposed to be associated with pS2pS5 diphosphorylated CTD in yeast (*48*). Considering pT4 and pS2 have 2/3 overlap in their interactomes (*13*), our results are consistent with the notion that pT4pS5 recruits TREX. Thus, we propose a model where TREX complex is recruited to pT4pS5 ‘double-letter’ CTD code over protein-coding genes at the early elongation stages and facilitates co-transcriptional splicing of mRNAs followed by their successful nucleocytoplasmic export (**Fig. 6E**). Over-representation of long transcripts among those that get deregulated upon T4 or TREX perturbation can be explained by similarity in both factors’ role in productive elongation and maintaining proper RNA Pol II transcriptional metrics, such as elongation speed in case of TREX and potentially, pT4. Given that genes with single T4 phosphorylation are not noticeably different from the average genes’ length, it is tempting to propose that pT4pS5 code is specifically ‘placed’ on longer genes to help them get fully processed by RNA Pol II with the help of TREX complex, which interacts with many other proteins inside the nuclear speckles (*49*).

One of potential study limitations is the inability to genetically engineer the double mutant cell line as Ser5 residue is necessary for cell survival and is considered ‘imperturbable’. However, since in our transcriptomic analyses we only focus on T4A-expressed genes simultaneously having pT4pS5 peaks, we believe that the filtered dataset indeed represents the direct effect of this combinatorial phosphomark. Another point is that our study is limited to exploration of pT4pS5 location and function in human cells, whereas pT4 and/or pT4pS5 functional assignment might be different in yeast cells. As an illustration, pT4 appears to selectively modulate transcription of specific class of non-coding snoRNA genes in yeast cells (*50*).

Thus, our results support the notion that pT4pS5 combinatorial phosphorylation marks on CTD function like “barcode” to selectively recruit transcription regulators for a specific cluster of genes. This barcode is recognized by regulators such as TREX and handled properly to achieve accurate transcription outcome. With known CTD-specific antibody masking effects and additional selective phosphatases, our experimental and analytical framework can potentially decipher other types of combinatorial CTD code. This data will further lead to functional annotation of each type of “barcode”, discovery of their similarities and differences, and eventually contribute to deeper mechanistic understanding of transcription in norm and disease.

## METHODS

### Cell culturing

Human embryonic kidney cells (HEK293/HEK293T) and osteosarcoma U2OS cells were purchased from ATCC (Manassas, VA). Cells were routinely cultured in Dulbecco’s-modified Eagle’s media (DMEM, Sigma-Aldrich, St. Louis, MO) supplemented with 10% Opti-Gold fetal bovine serum, FBS (GenDEPOT, Katy, TX) and 1% HyClone penicillin and streptomycin (P/S) mix (Cytiva, Marlborough, MA) at 37 °C in a humidified 5% CO_2_ atmosphere. Prior to the experiments, routinely cultured cell lines were confirmed mycoplasma free by the Mycoplasma qPCR kit (Minerva Biolabs, Skillman, NJ).

### Cloning

The yeast 26X T4E-CTD and mammalian 52X T4A/WT-CTD constructs were ordered as synthetic genes and cloned into a pET28a (Novogene, Sacramento, CA) derivative vector encoding a 6xHis-tag followed by a GST-tag and a 3C protease site or an N-terminal HA-tag mammalian expression vector. The 52X WT-CTD harboring α-amanitin resistance and an EYFP tag was from Addgene (plasmid #75284). Drosophila Ssu72 (1–195) and Symplekin (residues 19-351) were cloned into a pET28b vector encoding a 6x His-tag and SUMO tag. THOC2 shRNA (clone TRCN0000230704) was designed using Genetic Perturbation Platform from Broad Institute (www.portals.broadinstitute.org) and ordered as two DNA oligos from IDT DNA technologies (Coralville, IA). THOC2 shRNA was cloned into pLKO.5 vector using AgeI and EcoRI restriction enzymes and transformed into STBL3 *E.coli* strain (Amp+) (Invitrogen, #C737303). Non-mammalian shRNA control (pLKO.1-puro plasmid, #SHC002) was used as a shRNA control (Sigma-Aldrich).

### Transfections

To generate 52X T4A and 52X WT CTD HEK293T cells, transient transfection of 1.5 μg of either plasmid (per 1 well of 6-well plate) was performed using PEI (polyethylenimine) reagent (6.6 μl per 1 well). Next day after transfection, cells were selected with α-amanitin (2.5 μg/ml, Sigma-Aldrich, #A2263) for 72 hours with daily drug replenishment based on preliminary time-course experiment that had shown that untransfected cells are eliminated by this time-point.

To generate GFP-tagged Pol II-expressing U2OS cells for imaging, transient transfection of 1.5 μg of 52X WT CTD plasmid (per 1 well of 6-well plate) was performed using Fugene (Promega, Fitchburg, WI) following manufacturer’s instructions.

To generate shRNA-lentiviral stocks, HEK293T cells were seeded (0.5-1×10^6^ cells per 1 well of 6-well plate) and simultaneously transfected with a mixture of plasmids pLKO.5:gagpol:Vsvg (a kind gift from Pinglong Xu, Zhejiang University) with a ratio of 4:3:2 using PEI reagent, and the media was changed the next day. Lentivirus was collected at 24 h post-media change, filtered using 0.45 um filters and used within a week stored at +4 C. Target cells (HEK293) were infected with lentivirus in the presence of polybrene (final concentration 8 µg/ml) and spun down in plates for 30 min at 1,000 rpm. Selection with puromycin (Thermo Scientific, Rockford, IL) (final 1.5 µg/ml, for 2-3 days) was started when the cells became fully confluent, with untransfected cells as a control. Stable shRNA-expressing HEK293 cells were maintained in the complete media with 0.7-1 µg/ml of puromycin.

### Protein expression and purification

Symplekin (19-351), mutant Drosophila Ssu72 (C13D/D144N), and SUMO tagged Drosophila Ssu72 (1-195) were expressed in BL21 (DE3) cells and grown as single liter flasks at 37C in 50 ug/mL kanamycin LB or TB cultures to an OD_600_ of 0.4 to 0.9. Cultures were induced with 0.25 mM isopropyl-beta-D-thiogalactopyranoside (IPTG) overnight at 16C. Cultures were spun down, and the resultant pellets were resuspended in lysis buffer for sonication (5 cycles with 3 min intervals, 90 amp, 1 sec on, 5 sec off, on ice for 20-40 min process time) on a Sonic Ultrasonic Processor S-4000 (Misonix). Lysate was centrifuged for 30 to 45 minutes at 4C and 15,000 rpm and the supernatant was flowed through Ni^2+^/NTA beads (Qiagen, Germany). Beads were washed with wash buffer and eluted with elution buffer. Wild-type Ssu72, symplekin, and 3C protease were dialyzed together overnight, flowed through Ni^2+^/NTA beads, then loaded onto the FPLC. Mutant Ssu72 and symplekin were dialyzed together overnight, then loaded onto the FPLC, onto a Superdex 75 or 200. Symplekin and Ssu72 were combined in a 1:1.5 molar ratio, respectively, when dialyzed together. Dialysis and gel filtration used the same buffer, 25 mM Tris, 8.0, 100 mM NaCl, 10 mM BME.

### Dephosphorylation of the samples *in vitro*

Phosphatase (Ssu72, 50 µM) reactions were performed in a reaction buffer containing (20 mM MES, pH 6, 50 mM NaCl) for 30 min, at 28 C, 650 rpm on Thermomixer. The reaction time was optimized so that no further dephosphorylation occurred on the substrate.

### Dot blot

Serial dilutions of CTD peptides or Ssu72-treated CTD peptides were spotted (2 µl) on ready-to-use nitrocellulose membrane. The membrane was allowed to dry and blocked by soaking in TBS-T with 5% BSA for 30 min at room temperature. The membrane was then incubated with pT4 primary antibody (1:1,000, Active Motif, Carlsbad, CA) in TBS-T with 5% BSA for 45 min at room temperature. The membrane was washed three times with TBS-T for 5 min each. Then the membrane was incubated with anti-rat secondary antibody (1:10,000, LI-COR, Lincoln, NE) for 30 min at room temperature. The membrane was washed three times and visualized on LI-COR Odyssey CLx image reader.

### Western blot

Cells were lysed in RIPA lysis buffer (50mM Tris-Cl pH 8.0, 150 mM NaCl, NP-40, 0.5% sodium deoxycholate, 0.1% SDS) and 1× protease inhibitor cocktail (Roche, Indianapolis, IN). Protein concentrations were quantified with the Pierce BCA protein assay (Thermo Scientific). Typically, 20 μg of protein extracts were loaded and separated by SDS-PAGE gels. Blotting was performed with standard protocols using a PVDF membrane (Bio-Rad, Hercules, CA). Membranes were blocked for 1 h in blocking buffer (5% BSA in TBS-T) and probed with primary antibodies (rat pT4, 1:1,000, Active Motif, clone 6D7; rabbit THOC2, 1:500, Proteintech, Rosemont, IL, #55178-1-AP; mouse α-tubulin, 1:4,000, Sigma-Aldrich, #T6199; mouse histone H3 - only for nuclear/cytoplasmic lysates, 1:500, Sigma-Aldrich, clone 6.6.2) at 4 °C overnight. After three washes with TBS-T, the membranes were incubated with corresponding anti-rabbit, anti-mouse or anti-rat secondary IRDye 680RD antibody at 1:10,000 dilution (LI-COR) for 1 h at room temperature. After washing, membranes were visualized on LI-COR Odyssey CLx image reader.

### Immunofluorescence (IF) experiments

U2OS cells were transfected with 52X WT-CTD plasmids expressing GFP-tag using Fugene reagent (Promega) for 24 hours before harvest. Then, cells were fixed in 4% paraformaldehyde in PBS for 10 min at room temperature, permeabilized with PBS containing 0.1% Triton X-100 for 10 min at room temperature, then blocked in 2% bovine serum albumin (BSA) in PBS for 1 hour, and incubated with a primary antibody against THOC2 (Proteintech, #55178-1-AP, 1:50 dilution) overnight at 4°C. Next day, cells were stained with Alexa-labeled secondary antibody (Jackson, UK, #111-545-003, 1:500 dilution) for 1 hour at room temperature with extensive washing. Slides were stained with DAPI (Sigma-Aldrich, #MBD0015) and mounted with anti-Fade fluorescence mounting media (Abcam, Waltham, MA, #ab104135). Immunofluorescence images using DAPI, RFP, and GFP channels were obtained and analyzed using the Zeiss LSM710 confocal microscope and ImageJ software.

### MALDI-TOF experiments

108 µM 18-mer phosphopeptides and 5 µM Drosophila Ssu72/Symplekin (in 25 mM Tris, pH 8, 100 mM NaCl, 10 mM Beta-mercaptoethanol) were put into reaction buffer (20 mM MES, pH 6, 50 mM NaCl) and left for 30 min, at 28 C, 650 rpm on Thermomixer. Samples were adjusted to 0.1% trifluoroacetic acid (TFA), then desalted with Pierce C18 tips, 10 µl (Thermo Scientific, product #87782) according to manufacturer’s protocol with following changes: the wetting solution was 100% acetonitrile, the rinse solution was 0.1% TFA and did not include acetonitrile and the elution solution was 0.1% TFA, 50% acetonitrile. Elution solution included matrix (4mg/100 µl): 2,5 Dihydroxybenzoic acid (DHB) MALDI Matrix, Single Use, 3 x 8 microtube strips, 4 mg/microtube (Thermo Scientific, #90033). Samples were analyzed with FlexControl using Bruker autoflex maX MALDI-TOF/TOF. Analysis was conducted using flexAnalysis.

### Label-free proteomics sample preparation and CTD affinity purification

Erk2 (2.4 µM) was used to phosphorylate 1mg/ml of the 26x mutant yeast GST-CTD substrate where every Thr4 is mutate to a glutamate (T4E) in a 100 μL reaction for 1-24 hours. Likewise, a 26x yeast GST-T4E CTD substrate was incubated in a similar manner without any kinase treatment. Glutathione Agarose beads were washed in Buffer “C” (20 mM Tris pH 8.0, 150 mM NaCl, 10 mM BME) three times, and the treated GST-CTD samples were added to the beads and incubated overnight. A total of 200 million HEK293 cells were grown, collected, and the cell pellet was resuspended in Buffer “A” (10 mM HEPES pH 7.4, 100 mM NaCl, 300 mM Sucrose, 3 mM MgCl2, 0.5% Triton X-100, 1:100 Proteinase and Phosphatase Inhibitor). Cells were then vortexed, incubated on ice for 15 minutes, and centrifuged at 15,000 x g for 10 minutes at 4°C. The supernatant is discarded, and the cell pellet was resuspended in buffer “B” (10 mM Tris pH 8.0, 150 mM NaCl, 1:100 PPI) supplemented with 1:1000 benzonase. This mixture was incubated at room temperature for 1 hour and centrifuged at 15,000 x g for 10 minutes. The supernatant was collected as the nuclear fraction. After an overnight incubation, the GST-CTD bound beads were washed twice with buffer “C” and once with buffer “B”. The nuclear fraction was added to the substrate-bound beads and incubated at 4°C for 48 hours. Then the beads were centrifuged at 4,000 x g for 2 minutes at 4°C. The beads were washed twice with low salt buffer (20 mM Tris pH 8.0, 150 mM NaCl, 10% glycerol, 0.1% Triton X-100, and 1:100 PPI) and three times with high salt buffer (20 mM Tris pH 8.0, 500 mM NaCl, 10% glycerol, 0.1% Triton X-100, and 1:100 PPI). For each wash beads were incubated for 5 mins at 4°C. To the beads, 100 μl of elution buffer was added and the samples were placed on a rotator at 4°C for 2 hours. Then the beads were centrifuged at 4,000 x g for 2 minutes at 4°C, and the supernatant was collected for MS/MS analysis.

Pulldown samples were exchanged into 5 mM Tris-HCl using 3 kDa Amicon filters. Samples were then denatured in 2,2,2-trifluoroethanol (TFE) and 5 mM tris(2-carboxyethyl)phosphine (TCEP) at 55 °C for 45 min. Proteins were alkylated in the dark with 5.5 mM iodoacetamide, and the remaining iodoacetamide was quenched with 100 mM dithiothreitol (DTT). MS-grade trypsin was then added to the solution at an enzyme: protein ratio of 1:50, and the digestion reaction was incubated at 37 °C for 4 h. Trypsin was quenched by adding 10% formic acid, and the volume was reduced to 500 μL in a vacuum centrifuge. Samples were then filtered using a 10 kDa Amicon filter and desalted using Pierce C18 tips (Thermo Scientific). The samples were resuspended in 95% water, 5% acetonitrile, and 0.1% formic acid prior to MS.

### Proteomics mass spectrometry and protein identification

Peptides were separated on a 75 μm × 25 cm Acclaim PepMap100 C-18 column (Thermo Scientific) using a 5–50% acetonitrile + 0.1% formic acid gradient over 120 min and analyzed online by nanoelectrospray - ionization tandem MS on a Thermo Scientific Fusion Tribrid Orbitrap mass spectrometer, using a data-dependent acquisition strategy and analyzing two biological replicates per sample. Full precursor ion scans (MS1) were collected at high resolution (120,000). MS2 scans were acquired in the ion trap in rapid scan mode using the Top Speed acquisition method and fragmenting by collision-induced dissociation. Dynamic exclusion was activated with a 60 s exclusion time for ions selected more than once.

Proteins were identified with Proteome Discoverer 2.3 (Thermo Scientific), searching against the UniProt human reference proteome. Methionine oxidation [+15.995 Da], N-terminal acetylation [+42.011 Da], N-terminal methionine loss [−131.04 Da], and N-terminal methionine loss with the addition of acetylation [−89.03 Da] were all included as variable modifications. Peptides and proteins were identified using a 1% false discovery rate. To score changes in protein abundance between control and treated groups, we applied median centering followed by log2 transformation on a clean dataset, and then built linear models with limma in R (*29*). Enriched proteins were defined using a cut-off of log_2_FC > 0.6. Protein (PPI) networks were visualized using STRING database (v. 12.0) with built-in GO: Cellular component analysis accessed via ‘Analysis’ tab. Proteomic datasets were deposited under the accession number PXD059670.

### Co-Immunoprecipitation (Co-IP)

The 26x mutant (T4E) yeast GST-CTD samples with or without treatment by Erk2 kinase were prepared for co-IP in the exact same way as for label-free proteomics experiment (see before). After final bead centrifugation, nuclear supernatants were mixed with 2x Laemmli buffer and boiled for 5 min, then loaded (20 μL) and separated by SDS-PAGE gels. Blotting was performed with standard protocols using a PVDF membrane (Bio-Rad). Membranes were blocked for 1 h in blocking buffer (5% BSA in TBS-T) and probed with primary antibodies (mouse PRPF8, 1:500, Santa Cruz Biotechnology, Dallas, TX, #sc-55533; mouse His-tag, 1:500, Genscript, Piscataway, NJ, #A00186) at 4 °C overnight. After three washes with TBS-T, the membranes were incubated with corresponding anti-mouse secondary IRDye 680RD antibody at 1:10,000 dilution (LI-COR) for 1 h at room temperature. After washing, membranes were visualized on LI-COR Odyssey CLx image reader.

### RNA isolation, library preparation, and bulk and small RNA-Sequencing

Total RNA was isolated from HEK293T cells (at least ∼10^6^ cells per sample) using TRI reagent (Sigma-Aldrich) and DirectZol RNA Miniprep kit (Zymo Research, Irvine, CA, #R2050/#R2070). Nuclear and cytosolic cellular fractions for subsequent RNA extraction for qPCR experiments were separated using the following protocol. Cells were washed 2 times with 200 µl of ice-cold PBS and pelleted (3 min x 2,000 g). After resuspending in 200 µl of a buffer containing (10 mM HEPES KOH pH 7.4, 100 mM NaCl, 300 mM sucrose, 3 mM MgCl_2_, 0.5% Triton X-100), cells were vortexed and lysed on ice for 15 min. After centrifugation (10 min x 15,000 g at 4 C), supernatant was collected as cytosolic fraction. The pellet was washed with the same buffer, centrifuged again (1 min x 15,000 g at 4 C), and used as the nuclear fraction. Extraction of RNA from both fractions was performed using TRI Reagent.

Before sequencing, total RNA integrity was assessed by Novogene Co. using the RNA Nano 6000 assay kit of the Bioanalyzer 2100 system (Agilent Technologies, CA). Bulk RNA-Seq (after polyA enrichment, paired-end strategy 2X150bp) and small RNA-Seq (single-end strategy 50bp) libraries were prepared at Novogene Co. according to the manufacturer’s instructions for the Illumina platform (Illumina, San Diego, CA). The resulting libraries tagged with unique dual indices were checked for size and quality using the Agilent Bioanalyzer 2100. Libraries were loaded for sequencing on the NovaSeq 6000 instrument.

### Analyses of bulk RNA-seq data

Quality of raw reads was assessed using FastQC read quality reports (*51*). Adapter Illumina sequences and low-quality read ends were trimmed off by Trimmomatic v.0.38 with default parameters (*52*). Next, reads were aligned to human reference genome, GRCh38 version, using HISAT2 fast aligner v.2.2.1 with default parameters and -- unstranded type of library (*53*). Gencode v38 .gtf file was used as annotation gtf, including for genes’ lengths analysis. Lastly, mapped fragments were quantified by featureCounts v.2.0.1 in Galaxy (*54*). Differential expression was analyzed using DESeq2 v.1.30.1 in R; genes with FDR < 0.05 and a log_2_FC greater or less than 0.58 were considered as differentially expressed. Enrichment analysis of biological processes and cellular components was performed with ‘clusterProfiler’ v.3.18.1 R package (*55*), with genome wide annotation **‘**org.Hs.eg.db’ v.3.12.0 as gene universe. Alternative splicing analysis was performed using rMATS turbo v.4.1.2 (with parameters FDR < 0.05; ILD, inclusion level difference, ≥ 10%) (*42*). As input files for rMATS, alignment .bam files from HISAT2 mapper and gencode v38 annotation .gtf were used. JCEC files from rMATS output were further analyzed using custom scripts in R and SpliceTools (*44*) suite. Raw bulk RNA-seq data and processed counts were deposited in GEO under the accession number GSE262702.

Alternative splicing and differential gene expression upon TREX perturbation (THOC6 loss-of-function in neural progenitor cells) were re-analyzed using provided supplementary files (RSEM counts, JCEC files from rMATS pipeline) from GEO record GSE245121. In addition, differentially-expressed genes upon CHTOP knockdown in HEK293 cells were analyzed using salmon counts from another GEO record, GSE113952.

### Analyses of small RNA-seq data

Clean single-end data were trimmed with cutadapt v.4.6 (https://github.com/marcelm/cutadapt/blob/main/doc/guide.rst), using provided sequence ‘AGATCGGAAGAGCACACGTCT’ as 3’-adapter (-a) and sequence ‘GTTCAGAGTTCTACAGTCCGACGATC’ as 5’-adapter (-g). Only reads with minimal length of 17 bp were retained after trimming. Next, trimmed reads were mapped onto pre-indexed human reference genome (hg38) using bowtie2 v.2.3.5.1 (*56*) with the following parameters -N 0 -k 5 -- ignore-quals. Quantification was performed using featureCounts v.1.5.0 (*57*) with parameters ‘-t exon -O -s 1 -M - T 4’ and pre-filtered gencode hg38 .gtf file containing genomic coordinates of small RNAs: miRNAs, snoRNAs, snRNAs, and miscRNAs. Raw small RNA-seq data and processed counts were deposited in GEO under the accession number GSE286258.

### Nuclear/cytosolic fraction isolation for qPCR experiments

Cell pellets were washed twice with cold PBS, then resuspended in cold buffer A (10 mM HEPES pH 8.0, 5 mM MgCl_2_, 0.25 M sucrose, 0.1% NP-40, supplemented with 1x protease and phosphatase inhibitors and 1 mM DTT) and left on ice for 20 min. Next, cell lysates were centrifuged (10 min, 9,600 *g*, 4C), and supernatant was collected as cytosolic fraction. The intact nuclei (pellets) were resuspended in cold buffer B (25 mM HEPES pH 8.0, 20% glycerol, 1.5 mM MgCl_2_, 0.1 mM EDTA, 700 mM NaCl, supplemented with 1x protease and phosphatase inhibitors and 1 mM DTT), left 10 min on ice, and then sonicated using QSonica sonicator (Newtown, CT) (2 min, 3 sec on, 5 sec off, amp 65%). After sonication, nuclear lysates were centrifuged (15 min, 15,000 *g*, 4C) to get rid of insoluble material. Both fractions (nuclear and cytosolic) were immediately after isolation mixed with TRI reagent (Sigma-Aldrich) in 1:3 volume ratio, and then standard RNA isolation was performed.

### RT-qPCR

cDNA was generated using AzuraQuant cDNA synthesis kit (Azura Genomics, Raynham, MA) using manufacturer’s instructions. qPCR was done using the AzuraQuant Green Fast qPCR Mix Lo-Rox (Azura Genomics) in a ViiA-7 Real Time PCR system (Applied Biosystems). All qPCR experiments were conducted in biological triplicates. Relative gene expression was assessed using the ΔΔCt method normalized to GAPDH gene expression using technical duplicates. Specificity of amplification was checked with melting curves. All primers used in this study can be found in the **Supplementary Table S7.**

### Chromatin immunoprecipitation (ChIP) and ChIP-Sequencing

Briefly, single crosslinking of HEK293’s DNA was performed using 1% formaldehyde for 10 min. Crosslinking was quenched with 0.125 M glycine for 5 min. Cells were successively lysed in lysis buffer LB1 (50 mM HEPES-KOH, pH 7.5, 140 mM NaCl, 1 mM EDTA, 10% glycerol, 0.5% NP-40, 0.25% Triton X-100, 1× PI), LB2 (10 mM Tris-HCl, pH 8.0, 200 mM NaCl, 1 mM EDTA, 0.5 mM EGTA, 1× PI) and LB3 (10 mM Tris-HCl, pH 8.0, 100 mM NaCl, 1 mM EDTA, 0.5 mM EGTA, 0.1% Na-deoxycholate, 0.5% N-lauroylsarcosine, 1× PI). Chromatin was sonicated to an average size of ∼200–500 bp using UCD-200 Biorupter (30s on and 30 s off for 30 min). For pT4 IP, the active or catalytically dead (with mutation C13D/D144N) Ssu72 phosphatase was prepared in complex with symplekin, and then added (55 μM) to sonicated chromatin and incubated at 28°C for 30 min before immunoprecipitation. Next, a total of 5 μg of pT4 antibody (Active Motif, #61361) was pre-mixed in a 50 μl volume of Dynabeads protein G (Invitrogen) and was added to each sonicated chromatin sample and incubated overnight at 4°C. Input sample was processed in the same way, apart from the missing IP step. The chromatin-bound beads were washed two times with low salt buffer (0.1% Na Deoxycholate, 1% Triton X-100, 1mM EDTA, 50mM HEPES pH 7.5, 150mM NaCl), once with high salt wash buffer (0.1% Na Deoxycholate, 1% Triton X-100, 1mM EDTA, 50mM HEPES pH 7.5, 500mM NaCl), once with LiCl wash buffer (250mM LiCl, 0.5% NP-40, 0.5% Na-Deoxycholate, 1mM EDTA, 10mM Tris-Cl pH 8.0) and twice in TE buffer. The chromatin was reverse crosslinked overnight at 65°C with shaking at 750 rpm. After DNA extraction using phenol-chloroform, the DNA was resuspended in 10mM Tris-HCl pH 8.0. For sequencing, the extracted DNA was used to construct the ChIP-seq library using the NEBNext Ultra II DNA Library Prep Kit (single-end, x50bp) followed by sequencing with an Illumina HiSeq 3000.

### Analysis of ChIP-seq data

After initial assessment of read quality with fastQC, pT4 (treated with Ssu72 or Mutant) single-end ChIP-Seq reads were trimmed using TrimGalore! v.0.6.3 in Galaxy using default parameters, and then mapped onto human reference genome, hg19, using BWA v.0.7.17 (*58*) using default parameters. Alignments were filtered using samtools v.1.8 (mapq > 20), and then MACS2 v.2.2.7.1 in Galaxy (parameters: --broad; --broad-cutoff of q < 0.1) was used to call peaks for IP-samples against Input sample (*59*). Differential binding analysis was performed with DiffBind v.3.0.15 in R (*60, 61*) using three biological replicates of pT4 peaks (Condition: Ssu72 or Mutant) with batch effect correction (design = “∼Condition+Replicate”). Coverage tracks in .bigwig format and normalized either by Input read counts (log_2_Input) or RPKM were generated from filtered .bam files (mapq > 20) and visualized in IGV v.2.4.16 software. Bioconductor R package ‘chipseeker’ v.1.18.0 was used for differential peak annotation using gencode hg19 .gtf as a reference (*62*). For peak annotation, promoters were defined as (-1,000 bp, +1,000 bp from TSS) regions. TSS/TES profiling was done using ‘plotProfile’ or ‘plotHeatmap’ on matrices generated with 50-bp bins using the ‘computeMatrix’ function from the deeptools v.2.2.3 suite (*60*). Fingerprint plots for ChIP-Seq quality control were done using ‘plotFingerprint’, also from deeptools suite. Calculation of TSS, intragenic, and TES signals was performed using values derived with --outFileNameMatrix parameter of ‘computeMatrix’ function and custom R scripts. TSS and TES values were extracted at corresponding bins for every gene. Gene body signal was determined as the average signal across gene body bins (bins 40-80) for every gene. ChIP-seq data was added in GEO under the accession number GSE286362.

Flag-CHTOP ChIP-Seq data from GSE130992 record were re-analyzed using the following steps. After initial QC step, Illumina universal adapters were trimmed off using cutadapt v.4.6 with default parameters. Bowtie2 (v. 2.3.5.1) was used to align reads onto reference genome (hg38). After filtering out low-quality alignments (mapq < 10), MACS3 was used to call peaks (*p* < 0.001 determined by -cutoff-analysis parameter) in Flag-IP samples vs Input from the same GEO record. Bioconductor R package ‘chipseeker’ v.1.18.0 was used for peak annotation using gencode hg38 .gtf as a reference (*62*). For peak annotation, promoters were defined as (-1,000 bp, +1,000 bp from TSS) regions. As with pT4 ChIP-Seq, TSS/TES profiling (in RPKM) was done using matrices generated with 50-bp bins using the computeMatrix function from the deeptools v. 3.3.0.

RNA Pol II ChIP-Seq data (T4A vs Control and shTHOC5 vs Control) were accessed at GSE37519 and GSE173374, respectively. Supplementary processed files from the former record were directly used for deeptools visualization. For the latter record, we re-analyzed the data using the same data processing pipeline as for Flag-CHTOP ChIP-Seq. Coverage .bigwig files were prepared using --normalizeUsing CPM and --smoothLength 150 parameters. Next, Pol II metrics (namely, pausing index/PI) were calculated and visualized using a custom R pipeline, https://github.com/tailana703/PolII_metrics. Lastly, TT-Seq data [GSE173374] in THOC5-knockdown cells vs wild-type were visualized in IGV v.2.4.16.

### Structural modeling

A structure of Ssu72 in complex with a RNAPII CTD peptide phosphorylated at the Ser5 position (PDB Code: 4IMJ) was used as the starting point for our model. All residues in the CTD peptide were mutated in MAESTRO (version 13.5.128) so that a pT4 residue was positioned for dephosphorylation in the Ssu72 active site. The peptide was then energy-minimized in MAESTRO by applying the OPLS_2005 force field. The final model of pT4 in the active site of Ssu72 and the complex structure of pSer5 in the active site of Ssu72 were visualized in PyMOL (version 3.1.0a0 Open-Source).

### Statistical analyses

Satistical analyses were performed using RStudio v4.0.5 and GraphPad Prism v9. One-tailed or two-tailed t-test was used for comparing the two groups following normal distribution when applicable; in other cases, Wilcoxon rank sum test was used. Correlations were assessed using two-tailed Pearson *r* coefficients. Chi-squared test was used to compare distributions of categorical variables. Gene overlap analysis was performed using two-sided Fisher’s exact test in R. Protein bands and immunofluorecence images were quantified and compared using ImageJ software. *p* < 0.05 or FDR < 0.05 was considered as significant.

## Acknowledgments

This work was supported by the National Institutes of Health grant R01GM125882 (Y.J.Z.), R35GM148356 (Y.J.Z.), R35GM122480 (E.M.M.), Welch Foundation grant F-1515 (E.M.M.), and Army Research Office grant W911NF-12-1-0390 (E.M.M.). We also thank funding from L. Leon Campbell Professorship fund.

We are also grateful for technical assistance by the core facilities of the University of Texas at Austin. Computational analyses were performed using the Biomedical Research Computing Facility at UT Austin, Center for Biomedical Research Support, RRID# SCR_021979. Proteomic data was acquired in the UT Austin Center for Biomedical Research Support Biological Mass Spectrometry Facility, RRID# SCR_021728.

## Data and code availability

The data and code supporting the findings of this study are available from the corresponding author upon request. NGS data generated by this study have been deposited in GEO under the accession numbers: GSE286362 (ChIP-Seq), GSE286258 (RNA-Seq). Proteomic datasets can be accessed using accession number PXD059670.

