## Supplementary Figures for "Combinatorial phosphorylation on CTD of RNA polymerase II selectively controls transcription and export of protein-coding mRNAs"

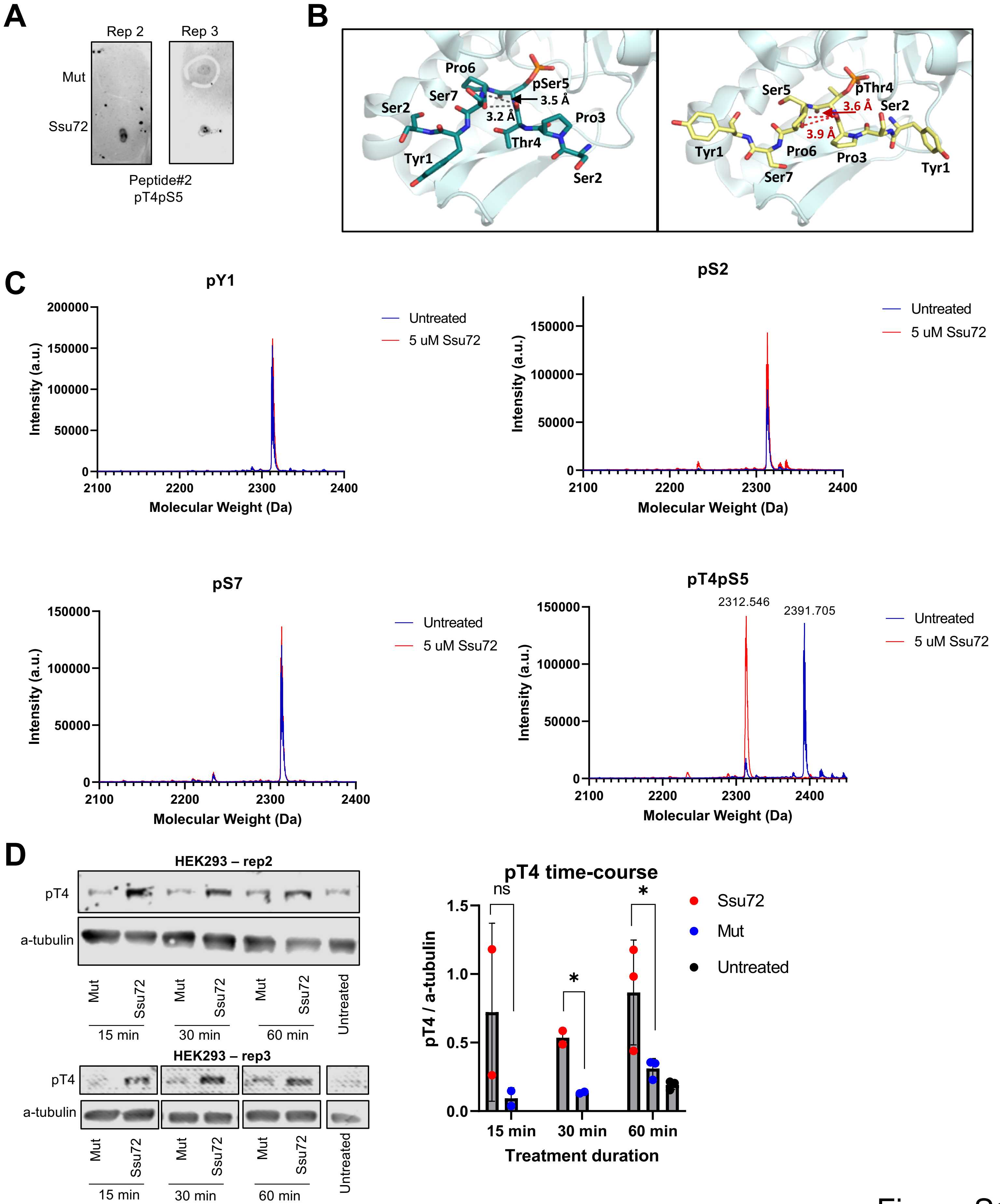
A
Rep 2
Rep 3
Mut
Ssu72
Peptide#2
pT4pS5
B
Pro6
pSer5
Ser7
3.5 Å
Ser2
3.2 Å
Pro3
Thr4
Tyr1
Ser2
Ser5
pThr4
3.6 Å
Ser2
3.9 Å
Tyr1
Pro6
Pro3
Ser7
Tyr1
C
pY1
pS2
pS7
pT4pS5
Untreated
5 uM Ssu72
Intensity (a.u.)
Molecular Weight (Da)
2312.546
2391.705
D
HEK293 – rep2
HEK293 – rep3
pT4
a-tubulin
Mut
Ssu72
Untreated
15 min
30 min
60 min
pT4 time-course
ns
*
pT4 / a-tubulin
Treatment duration
Ssu72
Mut
Untreated

Figure S1

**E**

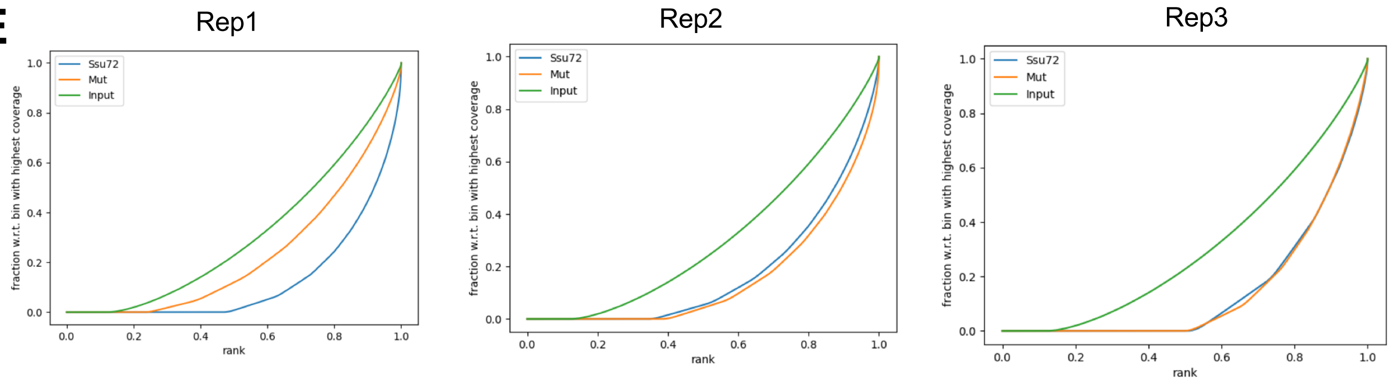


**F**

| Sample | Rep-licate | N, broad peaks ($q < 0.01$) | FRiP |
|---|---|---|---|
| pThr4-Ssu72 | 1 | 71,253 | 9% |
| pThr4-Ssu72 | 2 | 77,353 | 6% |
| pThr4-Ssu72 | 3 | 8,270 | 4% |
| pThr4-Mut | 1 | 9,281 | 4% |
| pThr4-Mut | 2 | 33,218 | 8% |
| pThr4-Mut | 3 | 4,287 | 3% |

**G**

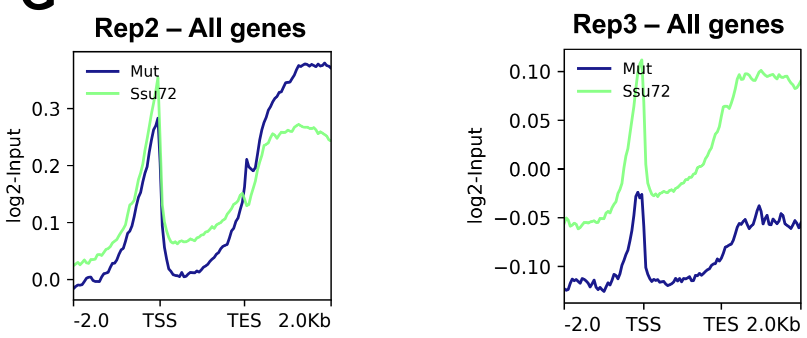


**H**

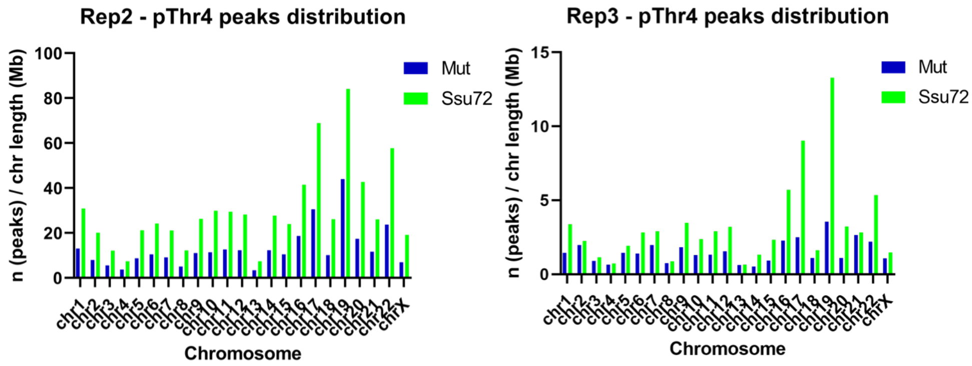


Figure S1

**Figure S1.** Validation of the framework to detect combinatorial CTD modifications using a pair of active and inactive phosphatases, Related to Figure 1.

**A**, Dot blot performed using pT4 antibody (6D7) and spotted doubly-phosphorylated peptide (pT4pS5) after incubation with active Ssu72 phosphatase or inactive phosphatase (Mut) for 30 min. Replicates 2-3 are shown.
**B**, Structure of pSer5 CTD peptide positioned in the active site of Ssu72 (*left*, PDB Code: 4IMJ) compared to a pT4 CTD peptide modelled into the active site of Ssu72 (*right*). Hydrogen bonds are shown as gray dashed lines, while disturbed bonds are shown as red dashed lines.
**C**, MALDI-TOF profiles of untreated (*in blue*) synthetic peptides having pY1, pS2, pS7, or double phosphomark pT4pS5 vs. corresponding peptides treated with 5 µM of active Ssu72 phosphatase (*in red*).
**D**, Time-course treatments (15, 30, 60 min) of HEK293 whole-cell lysates with active Ssu72 or its inactive counterpart (Mut). Shown are biological replicates 2-3 (*left*) and barplot with mean ± SD derived from 2-3 data points per condition (*right*). Comparison was performed using one-tailed paired *t*-test. $*p < 0.05$; ns - not significant.
**E**, Quality 'fingerprint' plots for pT4 ChIP-seq biological replicates (n = 3) built using 'deeptools'.
**F**, Quality characteristics of pT4 ChIP-seq biological replicates (n = 3) including numbers of called peaks and FRiP scores (fraction of reads in peaks).
**G**, Whole-genome metagene (n = 60,264) profile of Input-normalized pT4 ChIP-Seq signal under Ssu72 vs Mutant treatment. Region between TSS (transcription start site) and TES (transcription end site) is scaled to 2,000 bp. Every bin is 50 bp. Replicates 2-3 are shown.
**H,** pT4 peak distribution by chromosomes normalized by every chromosome's length in Mb upon Ssu72 vs Mutant treatment. Replicates 2-3 are shown.

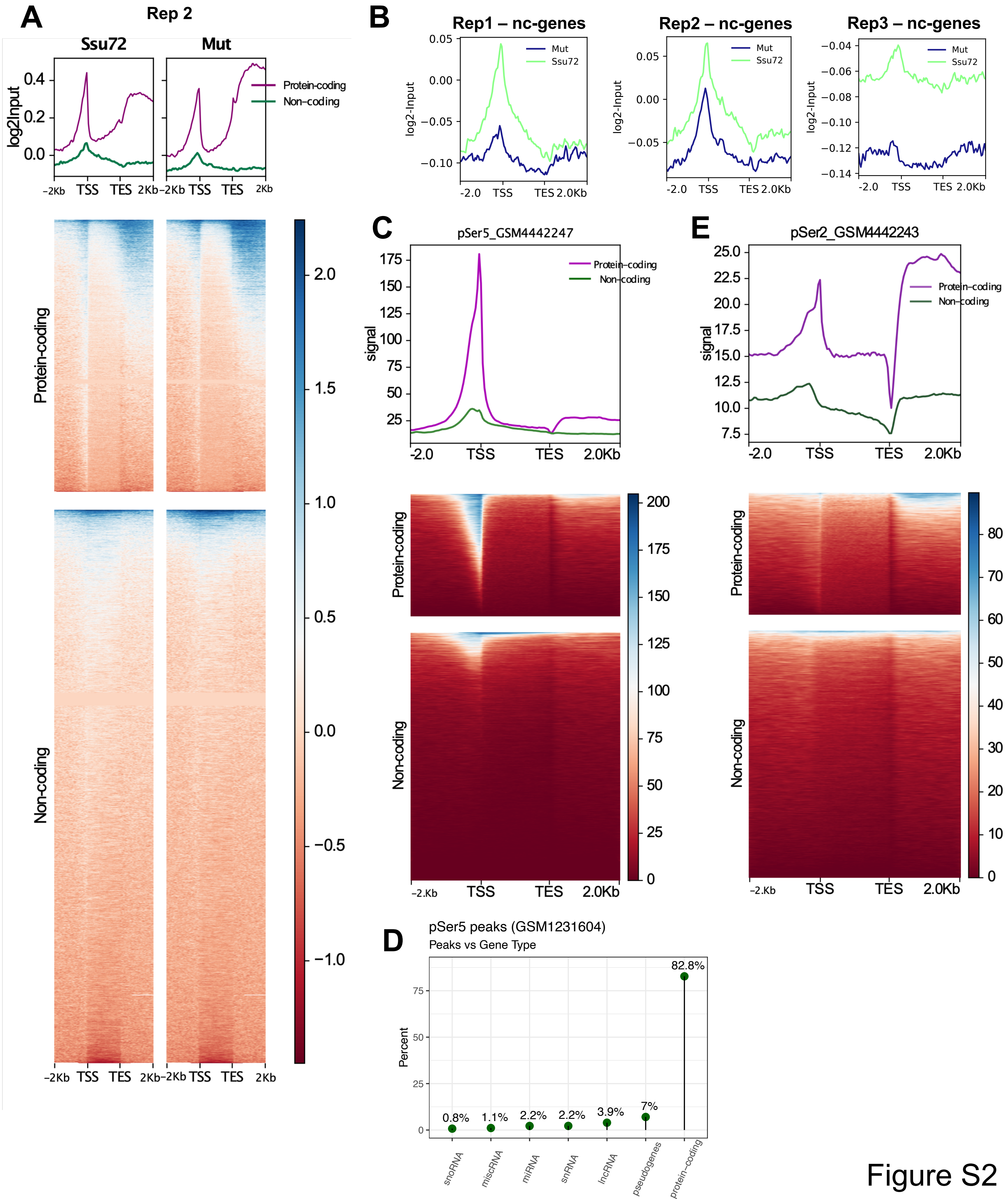

A
Rep 2
Ssu72
Mut
log2Input
Protein-coding
Non-coding
-2Kb TSS TES 2Kb
B
Rep1 – nc-genes
Rep2 – nc-genes
Rep3 – nc-genes
Mut
Ssu72
log2-Input
-2.0 TSS TES 2.0Kb
C
pSer5_GSM4442247
signal
Protein-coding
Non-coding
-2.0 TSS TES 2.0Kb
E
pSer2_GSM4442243
signal
Protein-coding
Non-coding
-2.0 TSS TES 2.0Kb
D
pSer5 peaks (GSM1231604)
Peaks vs Gene Type
Percent
0.8%
1.1%
2.2%
2.2%
3.9%
7%
82.8%
snoRNA
miscRNA
miRNA
snRNA
lncRNA
pseudogenes
protein-coding


Figure S2

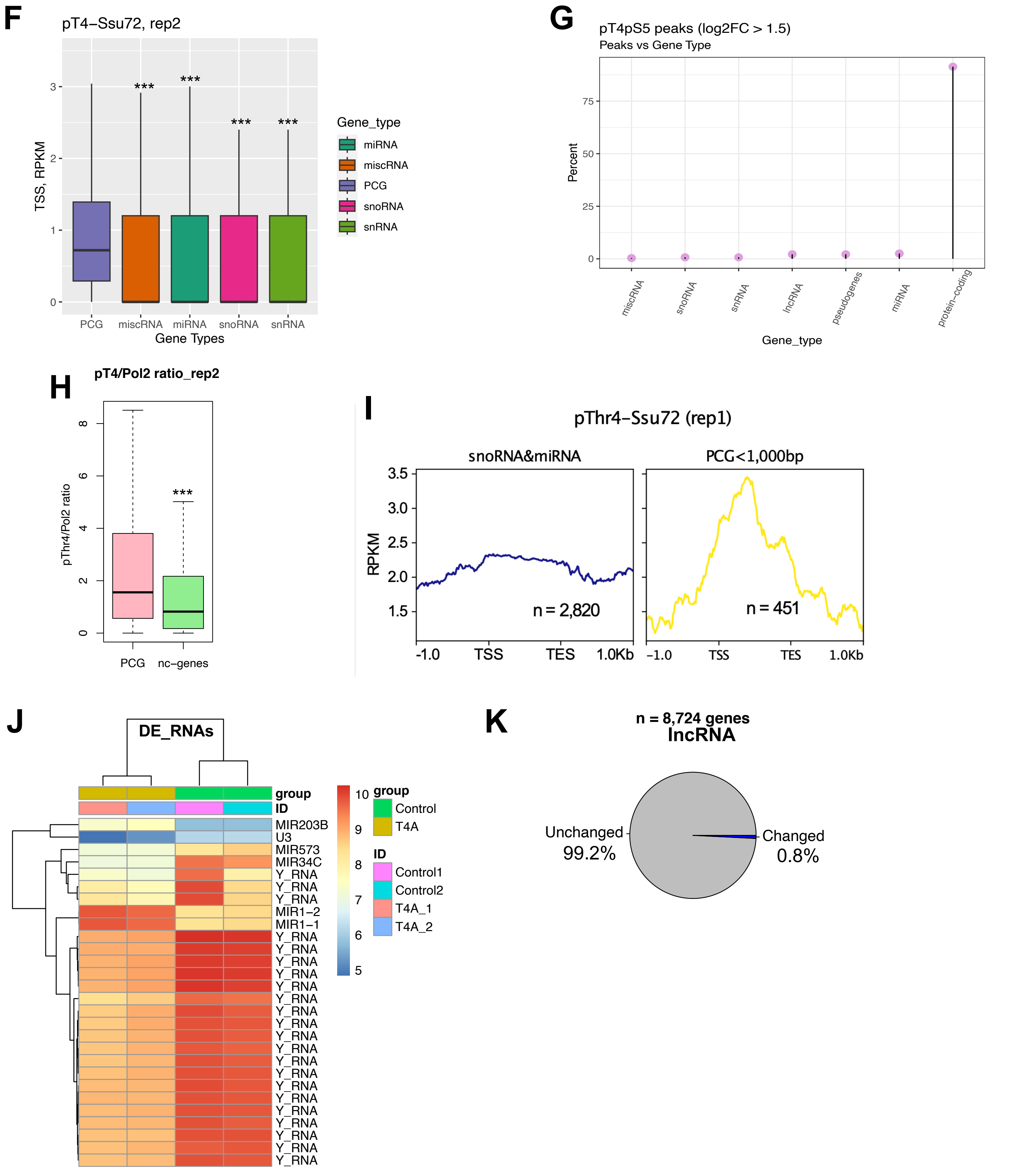
F
pT4−Ssu72, rep2
TSS, RPKM
0
1
2
3
***
***
***
***
PCG
miscRNA
miRNA
snoRNA
snRNA
Gene Types
Gene_type
miRNA
miscRNA
PCG
snoRNA
snRNA
G
pT4pS5 peaks (log2FC > 1.5)
Peaks vs Gene Type
Percent
0
25
50
75
miscRNA
snoRNA
snRNA
lncRNA
pseudogenes
miRNA
protein−coding
Gene_type
H
pT4/Pol2 ratio_rep2
pThr4/Pol2 ratio
0
2
4
6
8
***
PCG
nc−genes
I
pThr4−Ssu72 (rep1)
snoRNA&miRNA
PCG<1,000bp
RPKM
3.5
3.0
2.5
2.0
1.5
n = 2,820
n = 451
-1.0
TSS
TES
1.0Kb
J
DE_RNAs
group
ID
MIR203B
U3
MIR573
MIR34C
Y_RNA
Y_RNA
Y_RNA
MIR1−2
MIR1−1
Y_RNA
Y_RNA
Y_RNA
Y_RNA
Y_RNA
Y_RNA
Y_RNA
Y_RNA
Y_RNA
Y_RNA
Y_RNA
Y_RNA
Y_RNA
Y_RNA
Y_RNA
Y_RNA
Y_RNA
Y_RNA
Y_RNA
10
9
8
7
6
5
group
Control
T4A
ID
Control1
Control2
T4A_1
T4A_2
K
n = 8,724 genes
lncRNA
Unchanged
99.2%
Changed
0.8%

Figure S2

**Figure S2.** pT4pS5 phospho-sites are contained within protein-coding genes, Related to Figure 2.

**A**, Whole-genome metagene profile and heatmaps of Input-normalized pT4 ChIP-Seq signal under Ssu72 vs Mutant treatment on protein-coding (n = 19,838) and non-coding subsets (n = 40,426) separately. Region between TSS and TES is scaled to 2,000 bp. Every bin is 50 bp. Second replicate is shown.
**B**, Non-coding metagene (n = 40,426 genes) profile of Input-normalized pT4 ChIP-Seq signal under Ssu72 vs Mutant treatment (3 biological replicates are shown). Region between TSS and TES is scaled to 2,000 bp. Every bin is 50 bp. Heatmaps are sorted by mean value of each row (individual gene).
**C-E**, Whole-genome metagene profile and heatmaps of pSer5-CTD (**C**) and pSer2-CTD (**E**) ChIP-Seq signal on protein-coding (n = 19,838) and non-coding subsets (n = 40,426) separately. Region between TSS and TES is scaled to 2,000 bp. Every bin is 50 bp. Data was accessed using supplementary files at GEO (records GSM4442247 and GSM4442243 for pSer5 and pSer2, respectively). Heatmaps are sorted by mean value of each row (individual gene). **D**, Lollipop plot showing distribution of pSer5 peaks over different types of the genes. Data was accessed using supplementary files at GEO (record GSM1231604).
**F**, TSS signal (RPKM) of pT4 signal upon Ssu72 treatment in 5 gene subsets: protein-coding (PCG, n = 19,838), miscRNAs (n = 2,034), miRNAs (n = 3,055), snoRNAs (n = 1,457), snRNAs (n = 1,916) derived from human genome annotation (gencode, hg19). Statistical comparison was performed using Wilcoxon test, ***$p < 2.2e\text{-}16$. Second replicate is shown.
**G**, Lollipop chart of high-confidence pT4pS5 peaks ($log_2FC > 1.5$) distribution across different gene types.
**H**, Comparison of Pol2-normalized pT4 signal on protein-coding (PCG, n = 2,471) vs non-coding (nc, n = 934) genes with pT4pS5 peaks. Statistical comparison was performed using Wilcoxon test, ***$p < 2.2e\text{-}16$.
**I**, Analysis of pT4 signal upon Ssu72 unmasking (replicate 1) over sno&miRNA subset (n = 2,820) vs ultra-short protein-coding genes (1,000 bp or less, n = 451).
**J**, A heatmap of differentially-expressed small RNAs in T4A vs control cells. Most downregulated RNAs shown belong to Y-RNA class. Hierarchical clustering was performed using ‘pheatmap’ in R.
**K**, A pie chart showing relative number of deregulated lncRNAs (FDR < 0.05, both up- and down-regulated) in T4A vs WT cells (total n = 8,724).

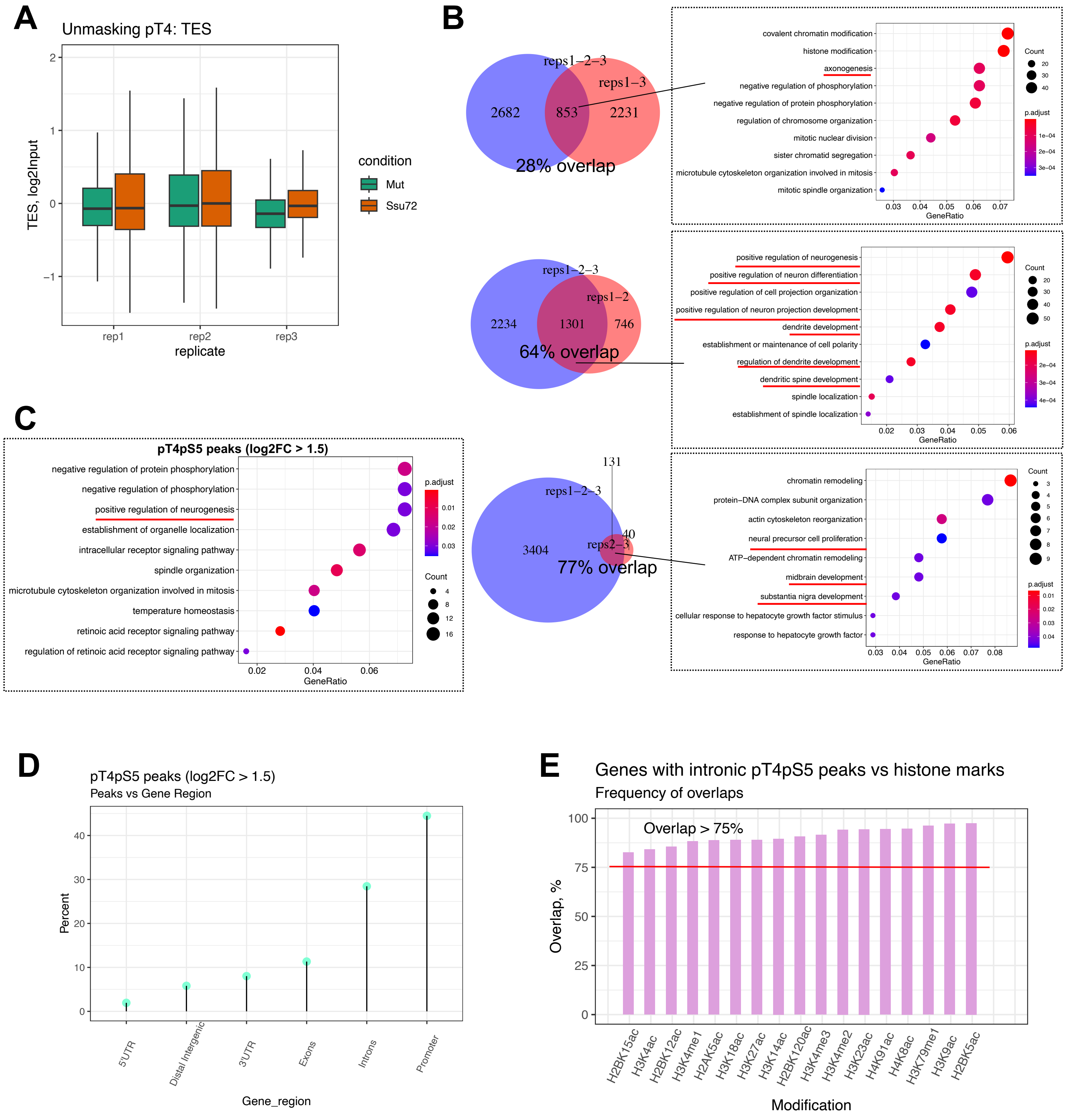
A
Unmasking pT4: TES
TES, log2Input
replicate
rep1
rep2
rep3
condition
Mut
Ssu72
B
reps1−2−3
reps1−3
2682
853
2231
28% overlap
covalent chromatin modification
histone modification
axonogenesis
negative regulation of phosphorylation
negative regulation of protein phosphorylation
regulation of chromosome organization
mitotic nuclear division
sister chromatid segregation
microtubule cytoskeleton organization involved in mitosis
mitotic spindle organization
GeneRatio
Count
p.adjust
reps1−2−3
reps1−2
2234
1301
746
64% overlap
positive regulation of neurogenesis
positive regulation of neuron differentiation
positive regulation of cell projection organization
positive regulation of neuron projection development
dendrite development
establishment or maintenance of cell polarity
regulation of dendrite development
dendritic spine development
spindle localization
establishment of spindle localization
GeneRatio
Count
p.adjust
131
reps1−2−3
3404
40
reps2−3
77% overlap
chromatin remodeling
protein−DNA complex subunit organization
actin cytoskeleton reorganization
neural precursor cell proliferation
ATP−dependent chromatin remodeling
midbrain development
substantia nigra development
cellular response to hepatocyte growth factor stimulus
response to hepatocyte growth factor
GeneRatio
Count
p.adjust
C
pT4pS5 peaks (log2FC > 1.5)
negative regulation of protein phosphorylation
negative regulation of phosphorylation
positive regulation of neurogenesis
establishment of organelle localization
intracellular receptor signaling pathway
spindle organization
microtubule cytoskeleton organization involved in mitosis
temperature homeostasis
retinoic acid receptor signaling pathway
regulation of retinoic acid receptor signaling pathway
GeneRatio
p.adjust
Count
D
pT4pS5 peaks (log2FC > 1.5)
Peaks vs Gene Region
Percent
5'UTR
Distal Intergenic
3'UTR
Exons
Introns
Promoter
Gene_region
E
Genes with intronic pT4pS5 peaks vs histone marks
Frequency of overlaps
Overlap > 75%
Overlap, %
H2BK15ac
H3K4ac
H2BK12ac
H3K4me1
H2AK5ac
H3K18ac
H3K27ac
H3K14ac
H2BK120ac
H3K4me3
H3K4me2
H3K23ac
H4K91ac
H4K8ac
H3K79me1
H3K9ac
H2BK5ac
Modification

Figure S3

**Figure S3.** pT4pS5 signal is enriched on protein-coding genes related to neurogenesis, Related to Figure 3.

**A**, Input-normalized pT4 signal at TES (transcription end site) on protein-coding genes (n = 19,838) upon treatment with either Ssu72 or Mutant in all performed replicates (n = 3).
**B**, Gene overlap analysis followed by GO: biological process analyses using 'DiffBind' and 'clusterProfiler' in R. Blue circle represents unique gene subset with called pT4pS5 peaks using all 3 performed paired replicates ("reps1-2-3"), whereas red circle represents unique gene subset with pT4pS5 peaks derived using iterations of duplicates ("reps1-3", "reps1-2", "reps2-3"). For each genetic overlap, GO:BP analysis identified enriched neurogenesis-related processes (highlighted in red, BH-corrected FDR < 0.05).
**C**, GO analysis of biological processes that were enriched among genes with high-confidence pT4pS5 peakset ($log_2FC > 1.5$). Analysis was performed using 'clusterProfiler' package in R and all human genes as gene universe, with BH-corrected FDR. Neuron-related terms are highlighted in red.
**D**, Lollipop chart of high-confidence pT4pS5 peaks ($log_2FC > 1.5$) distribution across different gene regions.
**E**, Frequency of overlaps (%overlap) between intronic pT4pS5 peaks and activating histone marks from Roadmap Epigenomic project. Neuronal cells were used as a query (H1-derived neuronal progenitor cells, "E007" sample).

A

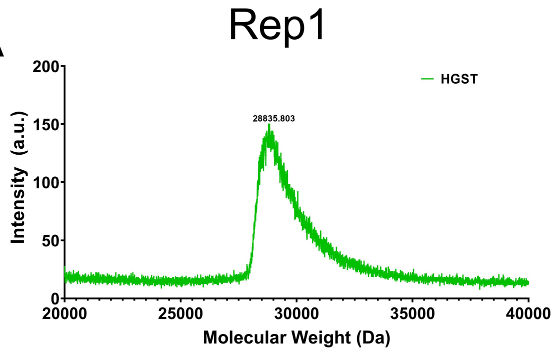


B

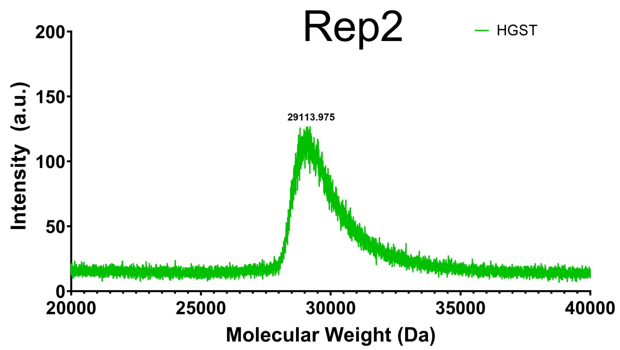


C

| Sample | Expected Mol. Wt. (Da) | Mol. Wt (Da) | # Phosphate Groups Added |
|---|---|---|---|
| HGST-26X CTD | 48,775.10 0 | 49,092.273 | - |
| HGST-26X CTD + Erk2 | - | 50,207.216 | ~14 |
| HGST-26X T4E (All) | 49,447.35 0 | 50,218.774 | - |
| HGST-26X T4E (All) + Erk2 | - | 51,341.232 | ~13 |
| HGST | 28,104.53 0 | 28,835.803 | - |

D

| Sample | Expected Mol. Wt. (Da) | Mol. Wt (Da) | # Phosphate Groups Added |
|---|---|---|---|
| HGST-26X CTD | 48,775.100 | 49,261.177 | - |
| HGST-26X CTD + Erk2 | - | 51,565.395 | ~29 |
| HGST-26X T4E (All) | 49,447.350 | 49,948.313 | - |
| HGST-26X T4E (All) + Erk2 | - | 52,202.257 | ~28 |
| HGST | 28,104.530 | 29,113.975 | - |

E

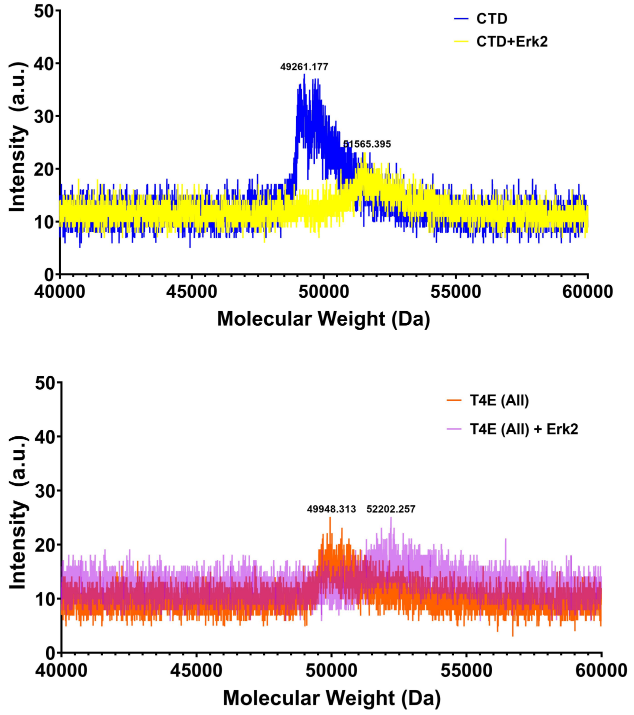


F

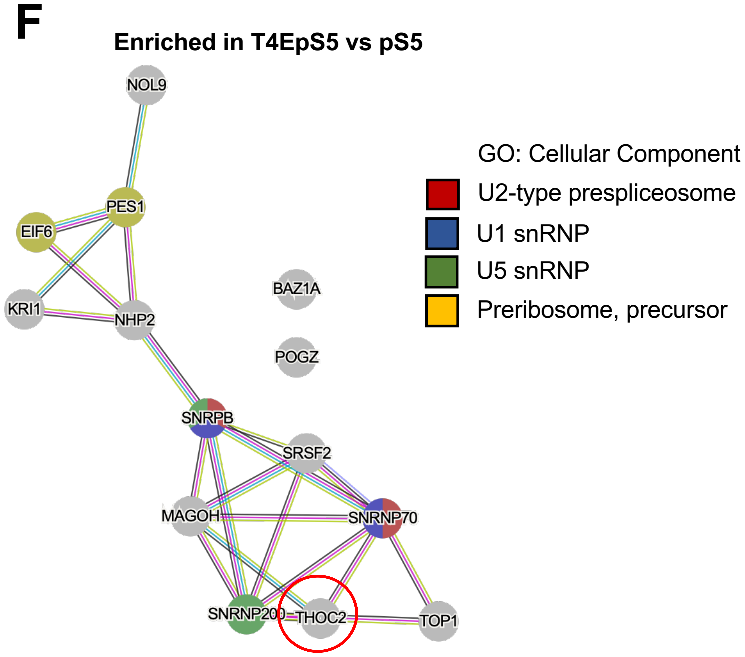


Figure S4

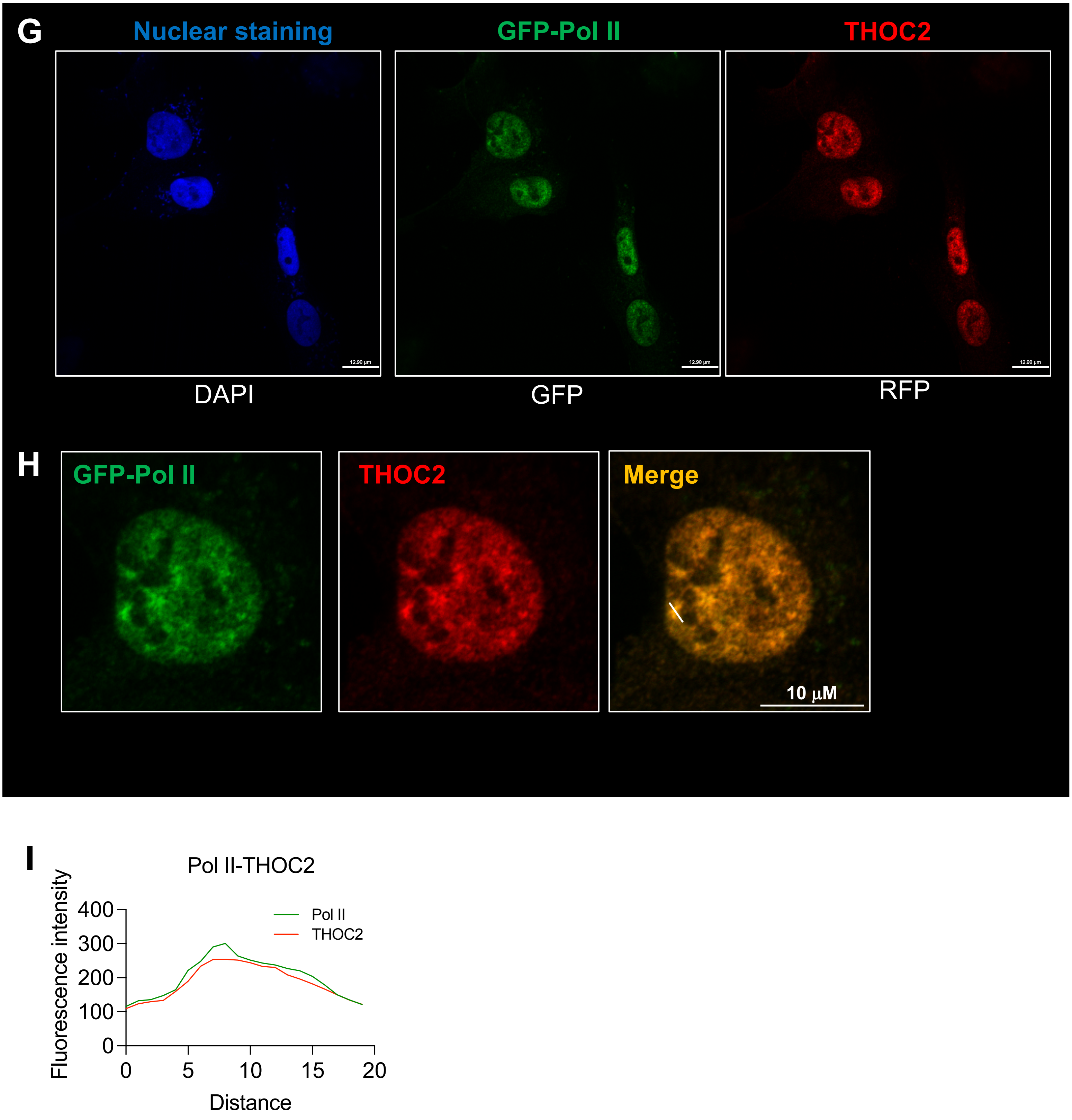
G
Nuclear staining
GFP-Pol II
THOC2
DAPI
GFP
RFP
H
GFP-Pol II
THOC2
Merge
10 μM
I
Pol II-THOC2
Fluorescence intensity
400
300
200
100
0
0
5
10
15
20
Distance
Pol II
THOC2

Figure S4

**Figure S4.** TREX complex is associated with pT4pS5 double marks, Related to Figure 4.

**A-B**, MALDI-TOF profiles of wild-type 26xCTD construct with human GST tag (HGST) for duplicates of proteomic pulldown experiments.
**C-D**, Calculation of numbers of phosphate groups added by Erk2 kinase (2.4 μM) to HGST-26x CTD and HGST-26x T4E constructs for duplicates of proteomic pulldown experiments.
**E**, MALDI-TOF profiles of wild-type 26xCTD construct (*upper*) and T4E-mutated 26xCTD construct (*lower*) before and after phosphorylation reaction using Erk2 kinase (2.4 μM). Second replicate is shown.
**F**, PPI (protein-protein interactions) network of proteins enriched in T4EpS5 pulldown vs single pS5 modification. Network was built using STRING databas; GO analysis (CC: cellular component) was performed using built-in 'Analysis' tab. THOC2 protein is highlighted.
**G-H**, IF images of GFP-Pol II (green channel, autofluorescence) expressing U2OS cells stained with DAPI (blue channel) and THOC2 antibody (red channel). Co-localization of GFP-Pol II and THOC2 is shown on merged image (yellow).
**I**, Co-localization of Pol II and THOC2 fluorescent signal across one of the well-formed puncta shown as white tick mark. Scale bar, 13 μm (**G**) or 10 μm (**H**).

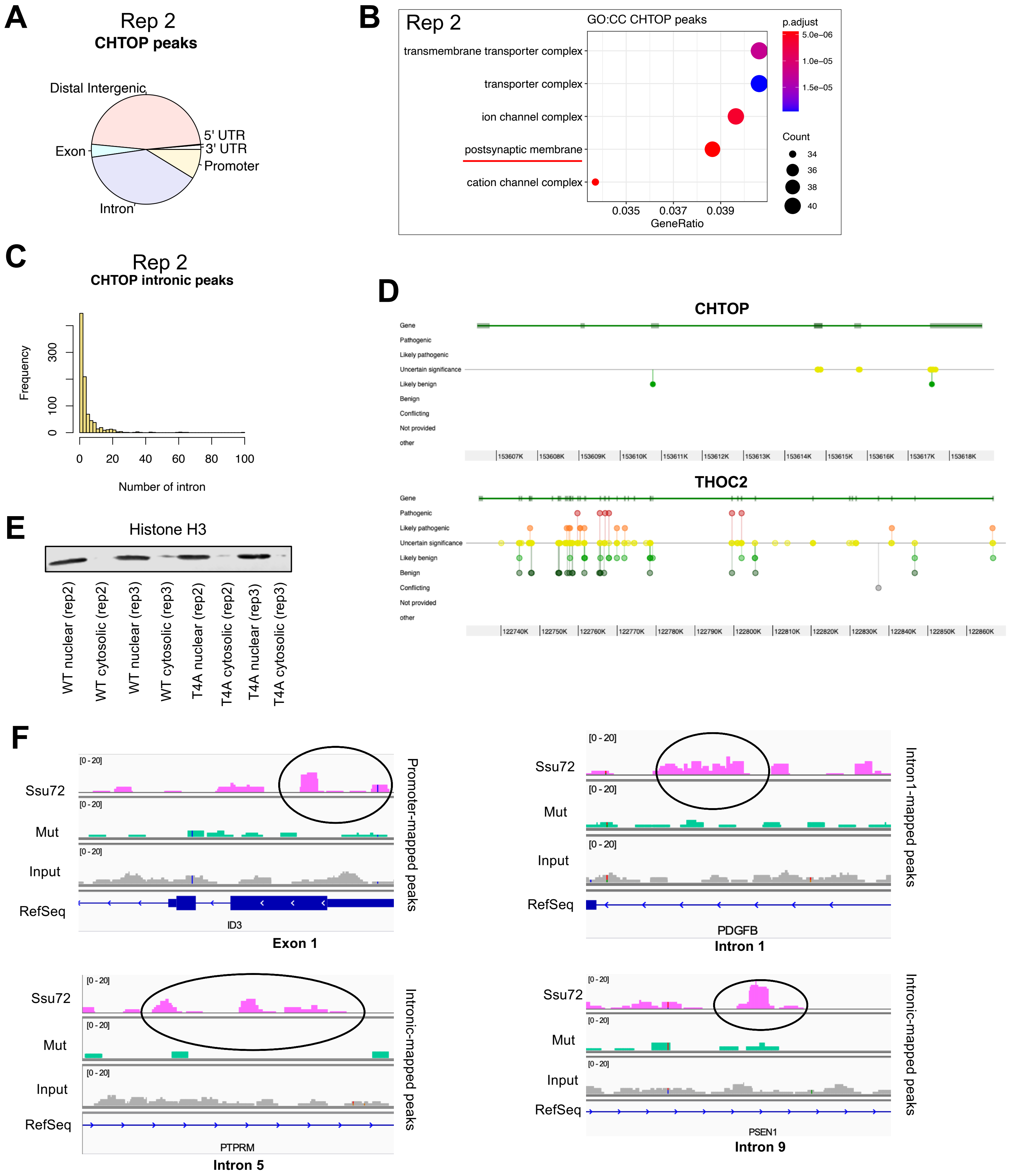
A
Rep 2
CHTOP peaks
Distal Intergenic
5' UTR
3' UTR
Exon
Promoter
Intron
B
Rep 2
GO:CC CHTOP peaks
transmembrane transporter complex
transporter complex
ion channel complex
postsynaptic membrane
cation channel complex
0.035
0.037
0.039
GeneRatio
p.adjust
5.0e−06
1.0e−05
1.5e−05
Count
34
36
38
40
C
Rep 2
CHTOP intronic peaks
Frequency
0
100
300
0
20
40
60
80
100
Number of intron
D
CHTOP
Gene
Pathogenic
Likely pathogenic
Uncertain significance
Likely benign
Benign
Conflicting
Not provided
other
153607K
153608K
153609K
153610K
153611K
153612K
153613K
153614K
153615K
153616K
153617K
153618K
THOC2
122740K
122750K
122760K
122770K
122780K
122790K
122800K
122810K
122820K
122830K
122840K
122850K
122860K
E
Histone H3
WT nuclear (rep2)
WT cytosolic (rep2)
WT nuclear (rep3)
WT cytosolic (rep3)
T4A nuclear (rep2)
T4A cytosolic (rep2)
T4A nuclear (rep3)
T4A cytosolic (rep3)
F
Ssu72
Mut
Input
RefSeq
[0 - 20]
ID3
Exon 1
Promoter-mapped peaks
PDGFB
Intron 1
Intron1-mapped peaks
PTPRM
Intron 5
Intronic-mapped peaks
PSEN1
Intron 9
Intronic-mapped peaks

Figure S5

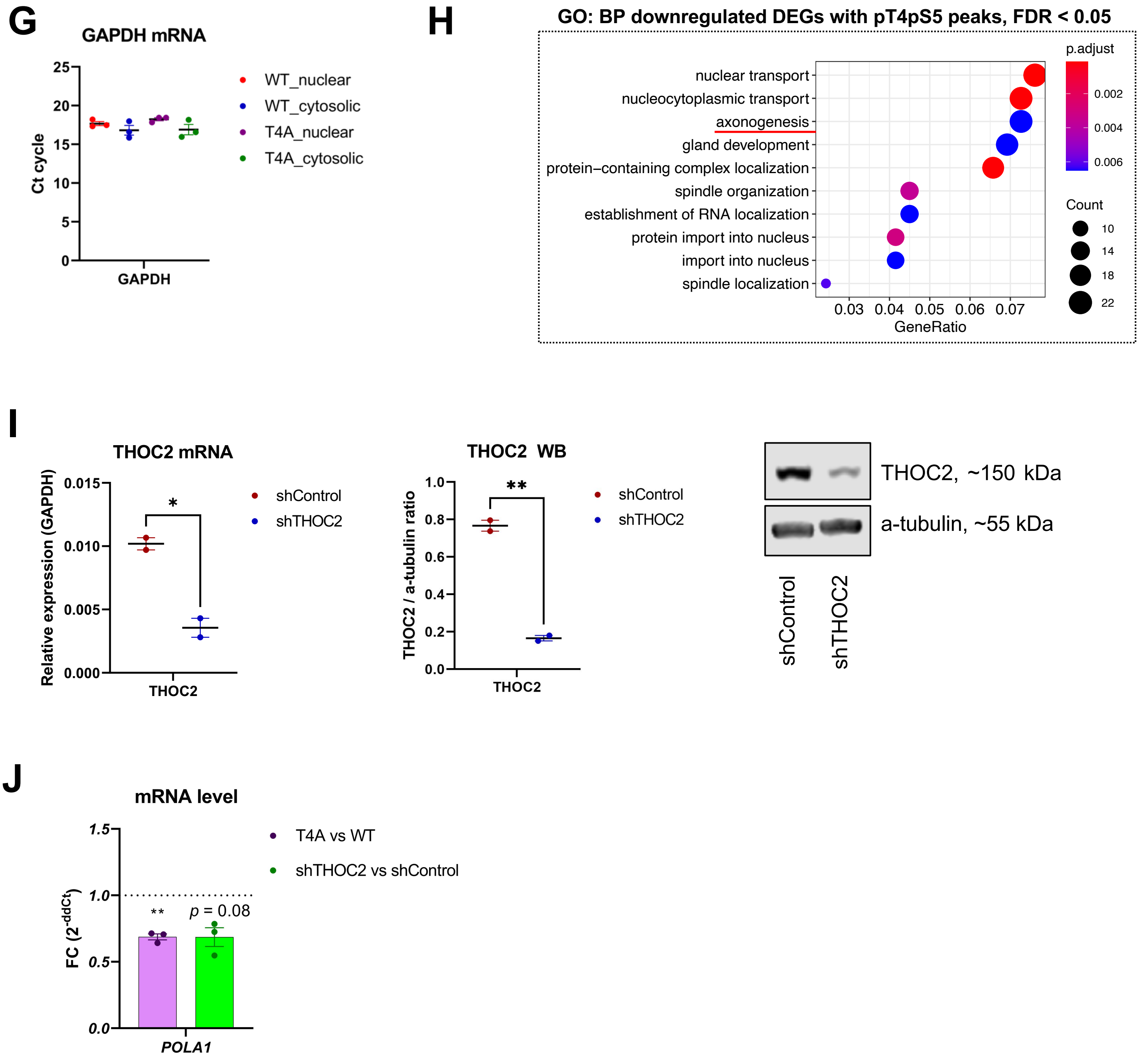
G
GAPDH mRNA
Ct cycle
25
20
15
10
5
0
GAPDH
WT_nuclear
WT_cytosolic
T4A_nuclear
T4A_cytosolic
H
GO: BP downregulated DEGs with pT4pS5 peaks, FDR < 0.05
nuclear transport
nucleocytoplasmic transport
axonogenesis
gland development
protein−containing complex localization
spindle organization
establishment of RNA localization
protein import into nucleus
import into nucleus
spindle localization
0.03
0.04
0.05
0.06
0.07
GeneRatio
p.adjust
0.002
0.004
0.006
Count
10
14
18
22
I
THOC2 mRNA
Relative expression (GAPDH)
0.015
0.010
0.005
0.000
*
THOC2
shControl
shTHOC2
THOC2 WB
THOC2 / a-tubulin ratio
1.0
0.8
0.6
0.4
0.2
0.0
**
THOC2
shControl
shTHOC2
THOC2, ~150 kDa
a-tubulin, ~55 kDa
shControl
shTHOC2
J
mRNA level
FC (2^-ddCt)
1.5
1.0
0.5
0.0
**
p = 0.08
POLA1
T4A vs WT
shTHOC2 vs shControl

Figure S5

**Figure S5.** Intronic-mapped pT4pS5 peaks recruit TREX for processing and nucleocytoplasmic export of mRNAs, Related to Figure 5.

**A**, Annotation of MACS3-called ($p < 0.001$) CHTOP peaks (GSE130992, replicate 2). Peak annotation was performed using 'chipseeker' R package.
**B**, GO analysis of cellular components enriched among genes with CHTOP peaks (GSE130992, replicate 2). Analysis was performed using 'clusterProfiler' package in R and all human genes as gene universe, with BH-corrected FDR.
**C**, Frequency histograms of CHTOP intronic peaks broke down by intron's number. Replicate 2 of CHTOP peaks (GSE130992) is shown.
**D**, Existence of pathogenic & likely pathogenic genetic variants, as well as variants-of-unknown-significance (VUS) within *THOC2* and *CHTOP* genes as illustrated in ClinVar database.
**E**, Histone H3 WB confirming optimal nuclear/cytosolic cellular fraction isolation for subsequent transcript export measurements with qPCR. Shown are biological replicates 2-3.
**F**, Exemplary IGV profiles of ChIP-Seq coverage in Ssu72, Mut, and Input samples. Shown are pT4pS5 peaks annotated to promoter region (*ID3*), intron 1 (*PDGFB*), and downstream introns (*PTPRM, PSEN1*).
**G**, Housekeeping *GAPDH* gene is expressed at similar levels across nuclear or cytosolic fractions. Shown are select 3 biological replicates.
**H**, GO analysis of biological processes enriched among downregulated genes with pT4pS5 peaks (FDR < 0.05). Analysis was performed using 'clusterProfiler' package in R and all human genes as gene universe, with BH-corrected FDR.
**I**, qPCR (*left*) and Western blotting (*middle*) validation of THOC2-knockdown in stably-generated shTHOC2 HEK293 cells. Shown are mean ± SEM derived from biological duplicates. Statistical comparison was performed using one-tailed paired *t*-test, *$p < 0.05$, **$p < 0.01$. *Right*, a representative WB of THOC2 and $\alpha$-tubulin in shTHOC2 and control cells.
**J**, Downregulation of *POLA1* gene expression upon T4 mutation (*in purple*) or THOC2-knockdown (*in green*). Shown are FC (mean ± SEM) calculated using ddCt method. Statistical comparison was performed using one-tailed paired *t*-test, **$p < 0.01$.

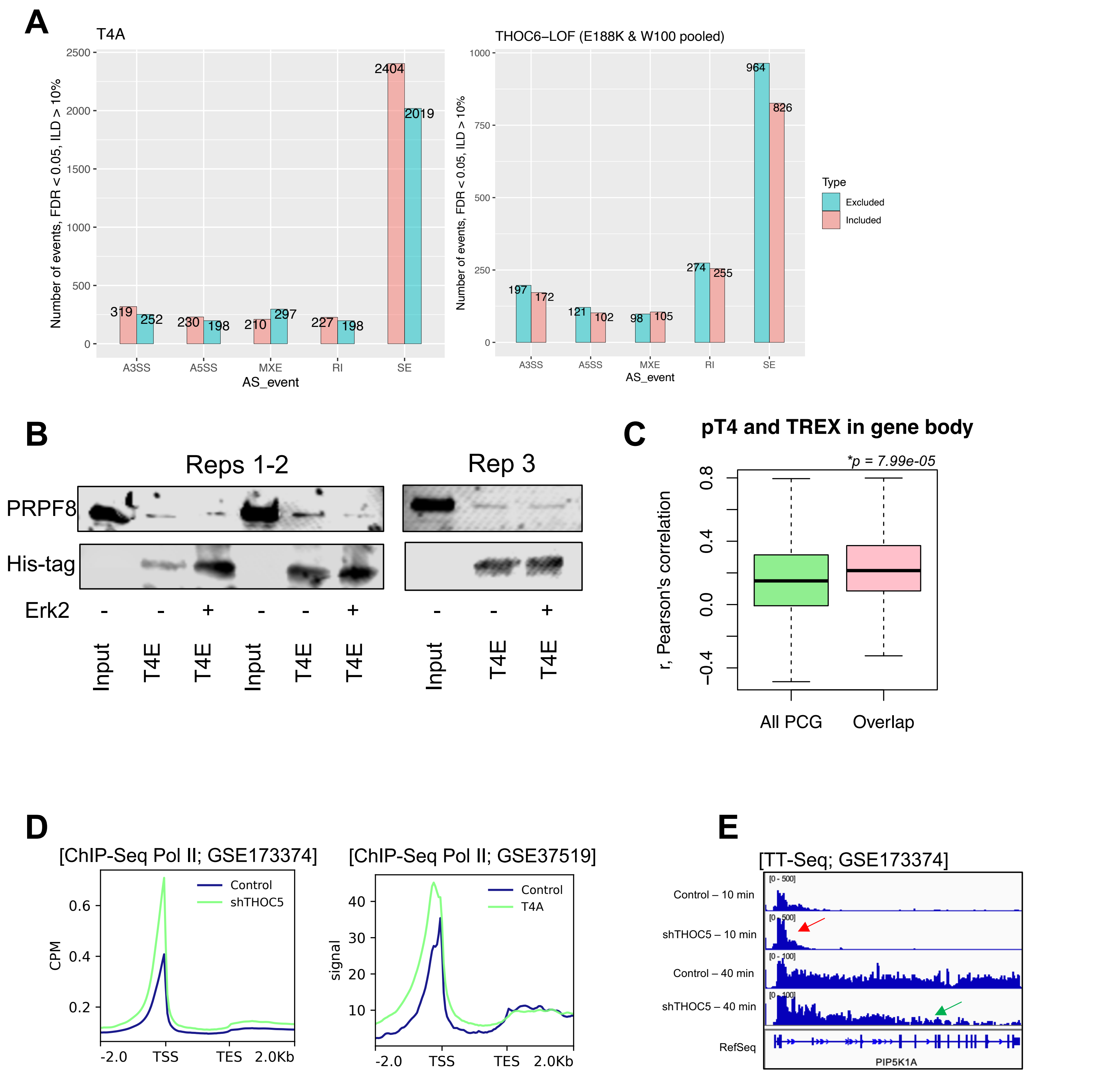
A
T4A
Number of events, FDR < 0.05, ILD > 10%
2500
2000
1500
1000
500
0
319
252
230
198
210
297
227
198
2404
2019
A3SS
A5SS
MXE
RI
SE
AS_event
THOC6–LOF (E188K & W100 pooled)
1000
750
500
250
197
172
121
102
98
105
274
255
964
826
Type
Excluded
Included
B
Reps 1-2
Rep 3
PRPF8
His-tag
Erk2
Input
T4E
C
pT4 and TREX in gene body
*p = 7.99e-05
r, Pearson's correlation
0.8
0.4
0.0
−0.4
All PCG
Overlap
D
[ChIP-Seq Pol II; GSE173374]
CPM
0.6
0.4
0.2
Control
shTHOC5
-2.0
TSS
TES
2.0Kb
[ChIP-Seq Pol II; GSE37519]
signal
40
30
20
10
Control
T4A
E
[TT-Seq; GSE173374]
Control – 10 min
shTHOC5 – 10 min
Control – 40 min
shTHOC5 – 40 min
RefSeq
[0 - 500]
[0 - 100]
PIP5K1A

Figure S6

F

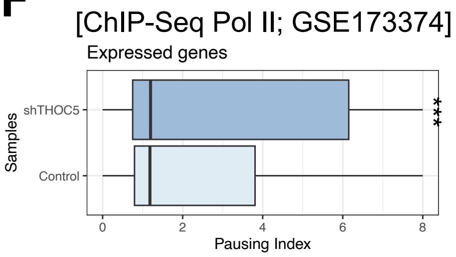


G

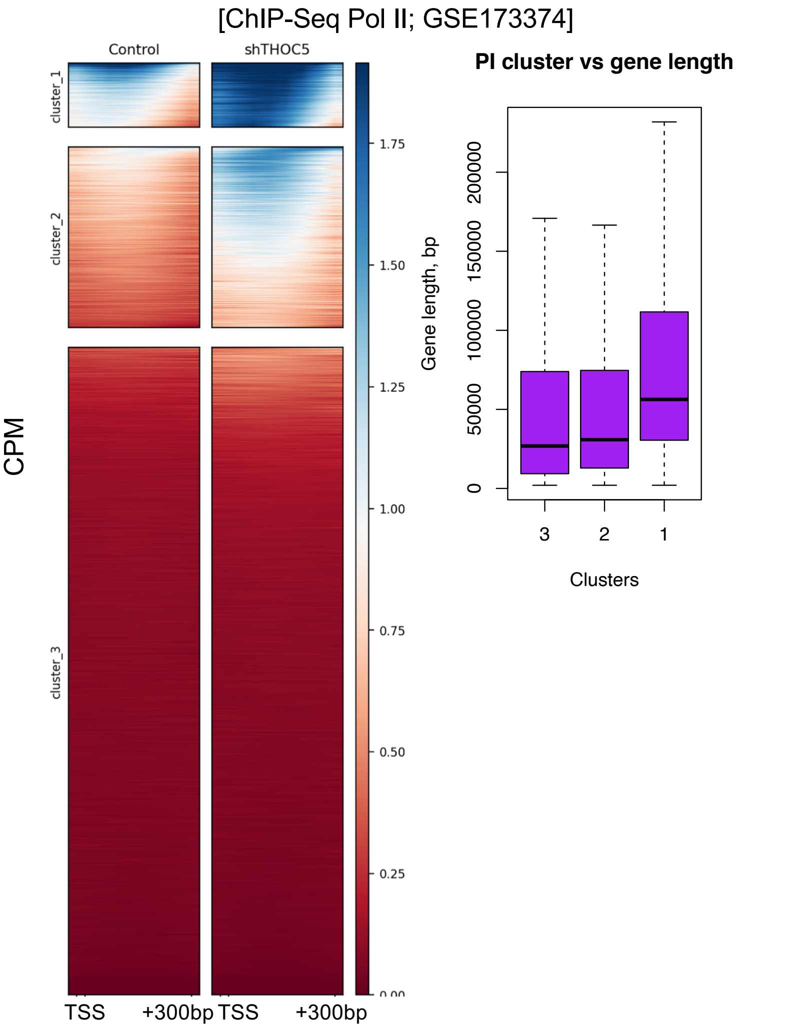


H

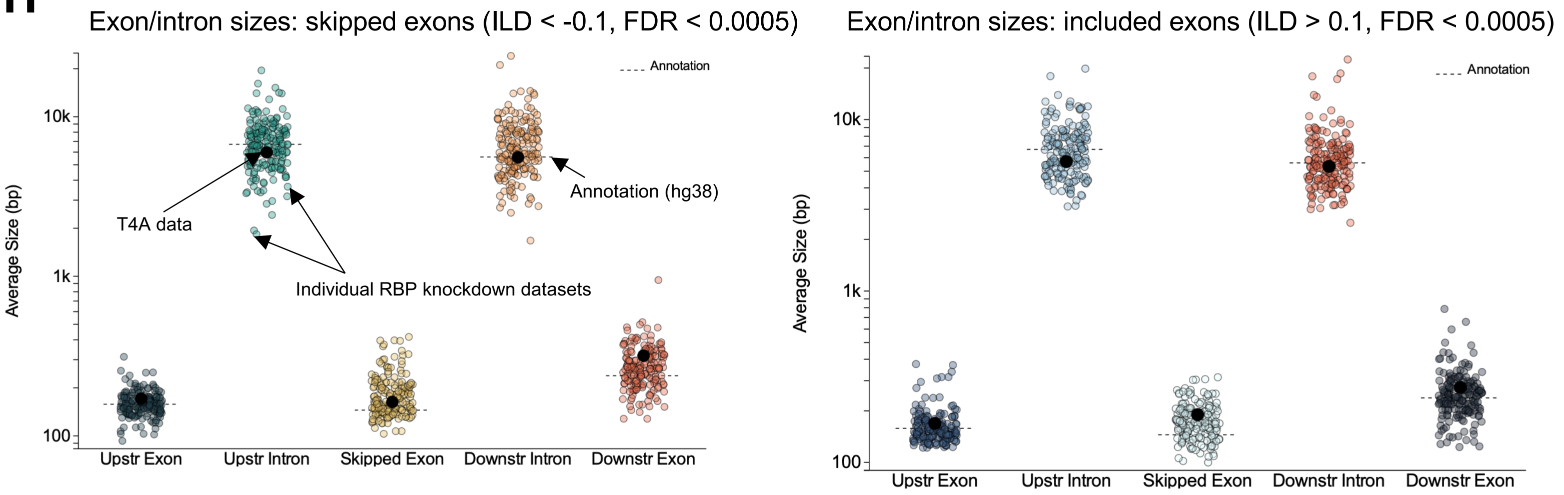


Figure S6

**Figure S6.** The pT4pS5 marks drive splicing of lengthy neurogenesis-related genes through TREX, Related to Figure 6.

**A**, Absolute numbers of alternative splicing events (FDR < 0.05, ILD > 10%) in T4A vs WT HEK293 cells (*left*) and THOC6 loss-of-function neural cells vs control (GSE245121, *right*). Shown are events of 5 types detected by rMATS pipeline: A3SS – alternative 3'-splice sites, A5SS – alternative 5'-splice sites, MXE – mutually exclusive exons, RI – retained introns, SE – skipped exons. Included events, ILD > 0; excluded events, ILD < 0.
**B**, Co-IP experiments (n = 3 biological replicates) confirming interaction of PRPF8 with singly-modified 26x-T4E and Erk2-derived 26x-T4EpS5 'baits'.
**C**, Correlation of binned ChIP-Seq signal of pT4 (upon Ssu72 treatment, replicate 1) and CHTOP (GSE130992) on all protein-coding genes (n = 19,840, in *green*) vs genetic overlap from **Fig. 6C** (n = 158 genes, *in pink*). Statistical comparison was performed using Wilcoxon test.
**D**, ChIP-Seq Pol II profiles on protein-coding genes (n = 19,840) in THOC5-knockdown cells vs Control (*left*, GSE173374), and T4A cells vs Control (*right*, GSE37519). Region between TSS and TES is scaled to 2,000 bp. Every bin is 50 bp.
**E**, TT-Seq data (GSE173374) visualization on a representative long gene with detected pT4pS5 double phosphorylation – *PIP5K1A*. THOC5-knockdown resulted in slower elongation speed and delayed transcription wave.
**F**, Pausing index (PI) calculation of Pol II in shTHOC5 cells vs control cells (GSE173374). Shown are average PI values derived from duplicate experiments. Statistical comparison was performed using paired Wilcoxon test, ***$p$ < 2.2e-16.
**G**, Heatmaps of Pol II occupancy on protein-coding genes derived after unsupervised k-means clustering of ChIP-Seq Pol II signal (CPM) in shTHOC5 vs control cells (GSE173374). Shown is (TSS; TSS+300 bp) region. Cluster 1 with the highest Pol II storage on the promoter represents especially lengthy genes (median length > 50 kb).
**H**, Average sizes of skipped (ILD < -0.1, FDR < 0.0005, *left*) and included (ILD > 0.1, FDR < 0.0005, *right*) exons along with corresponding upstream/downstream introns and exons' sizes upon T4A mutation (shown as thick black dot) in comparison with human annotation data (dashed line) and individual RNA-binding protein knockdown datasets (all other dots). Analysis was performed using SpliceTools suite.
